# Habitat-specificity in SAR11 is associated with a handful of genes under high selection

**DOI:** 10.1101/2024.12.23.630198

**Authors:** Sarah J. Tucker, Kelle C. Freel, A. Murat Eren, Michael S. Rappé

## Abstract

The order *Pelagibacterales* (SAR11) is the most abundant group of heterotrophic bacteria in the global surface ocean, where individual sublineages likely play distinct roles in oceanic biogeochemical cycles. Yet, understanding the determinants of niche partitioning within SAR11 has been a formidable challenge due to the high genetic diversity within individual SAR11 sublineages and the limited availability of high-quality genomes from both cultivation and metagenomic reconstruction. Here, we take advantage of 71 new SAR11 genomes from strains we isolated from the tropical Pacific Ocean to evaluate the distribution of metabolic traits across the *Pelagibacteraceae,* a recently classified family within the order *Pelagibacterales* encompassing subgroups Ia and Ib. Our analyses of metagenomes generated from stations where the strains were isolated reveals distinct habitat preferences across SAR11 genera for coastal or offshore environments, and subtle but systematic differences in metabolic potential that support these observations. We also observe higher levels of selective forces acting on habitat-specific metabolic genes linked to SAR11 fitness and polyphyletic distributions of habitat preferences and metabolic traits across SAR11 genera, suggesting that contrasting lifestyles have emerged across multiple lineages independently. Together, these insights reveal niche-partitioning within sympatric and parapatric populations of SAR11 and demonstrate that the immense genomic diversity of SAR11 bacteria naturally segregates into ecologically and genetically cohesive units, or ecotypes, that vary in spatial distributions in the tropical Pacific.

## Main

Bacterial populations typically harbor extensive variation in gene content that can be difficult to delineate into discrete ecological or evolutionary units, despite the significant impact such differences may have on the biogeochemical roles, ecological interactions, and fitness of these microorganisms (Louca et al. 2016; Hao et al. 2020; Wang et al. 2023). Comparative genomics has enabled the identification of gene content variation that underpins distinct ecologies and contributes to the process of niche diversification (Cordero and Polz 2014; Sheppard et al. 2018; Whelan et al. 2021). However, because these analyses most often use a collection of genomes isolated from diverse environments where genes have evolved under a range of selective pressures (Conrad et al. 2022), associating variations in gene content with specific forces of selection and identifying crucial genes that likely serve as determinants of fitness under specific conditions remains a formidable challenge.

The bacterial order *Pelagibacterales* (SAR11) is one of the most abundant and diverse bacterial groups in the surface oceans (Morris et al. 2002; Brown et al. 2012; Grote et al. 2012), where even a single SAR11 population can exhibit substantial sequence diversity in the environment (Delmont et al. 2019). Defining the genomic variation underlying distinct spatial distributions of SAR11 populations and how gene content differs across environmental conditions is vital to further understanding ocean biogeochemical cycling, eco-evolutionary relationships, and the biology of a ubiquitous and dominant lineage in the surface oceans. This task remains difficult because assessing the loss or gain of ecologically-relevant genes or metabolic pathways in SAR11 necessitates a robust collection of high quality genome sequences that represent a spectrum of both ecological and genomic diversity. SAR11 cells are historically recalcitrant to laboratory cultivation, and as such only a small portion of the total diversity of naturally occurring SAR11 populations can be interrogated via high quality genomes such as those from isolated strains (Haro-Moreno et al. 2020). Yet even with a small set of isolate genomes, SAR11 has become a key model to help elucidate fundamental processes of ecology and evolution such as genome streamlining (Giovannoni et al. 2005), marine oligotrophy (Noell et al. 2023), co-evolution (Morris et al. 2012; Braakman et al. 2017), drivers of genetic diversity and evolution within organisms of large population sizes (Vergin et al. 2007; Delmont et al. 2019), structure-aware investigations of microbial population genetics (Kiefl et al. 2023), and ocean biogeochemical cycling (Grant et al. 2019; White et al. 2019; Acker et al. 2022).

In our companion work (Freel et al. 2024), we address the gap in the available genomic resources with novel SAR11 isolates that originate from a geographically constrained environment within the tropical Pacific, and develop a phylogenomic backbone that unites evolution with ecology to understand this diverse clade. Freel et al., (2024) delineated four families within SAR11, including the family *Pelagibacteraceae* which encompasses the historical SAR11 subgroups Ia and Ib. Here, we illuminate metabolic diversity found within *Pelagibacteraceae* and identify the determinants of niche differentiation within co-occurring *Pelagibacteraceae* populations by combining 71 of the new isolates with 21 previously sequenced *Pelagibacteraceae* isolate genomes. By taking advantage of metagenomes from the prominent source of isolation, we apply an integrated ‘omics analysis to unveil differences in *Pelagibacteraceae* metabolic features across coastal and offshore habitats and identify metabolic genes under high selective pressures.

## Results

### Ecology and evolution of *Pelagibacteraceae* reveals strong habitat-specificity

By analyzing 92 high quality genomes from isolates obtained predominantly from coastal Kāneʻohe Bay, Oʻahu, Hawaiʻi, and the adjacent offshore (**Fig. 1, Supplementary file 1a**), we reveal that the genomic diversity of cultured *Pelagibacteraceae* partitions into eleven discrete monophyletic clusters, seven of which include isolates from the Kāneʻohe Bay Time-series (KByT; **Fig. 1, Extended Data Fig. 1a**). The 92 genomes belong to both previously-described and new clades, which we recently described as eleven genera (Freel et al. 2024; **Fig. 1**, **Supplementary file 1a**). The within-genus genome-wide average nucleotide identity (gANI) differed among genera, with the lowest within-genus average gANI found in *Proprepelagibacter* (87.1±0.0%, mean ± sd) and the highest within-genus average gANI found in *Littoralipelagibacter* (95.8±0.0; **Supplementary file 1b**). The average of all within-genus genome-wide average nucleotide identities was 92.3±2.7%.

**Fig. 1.**
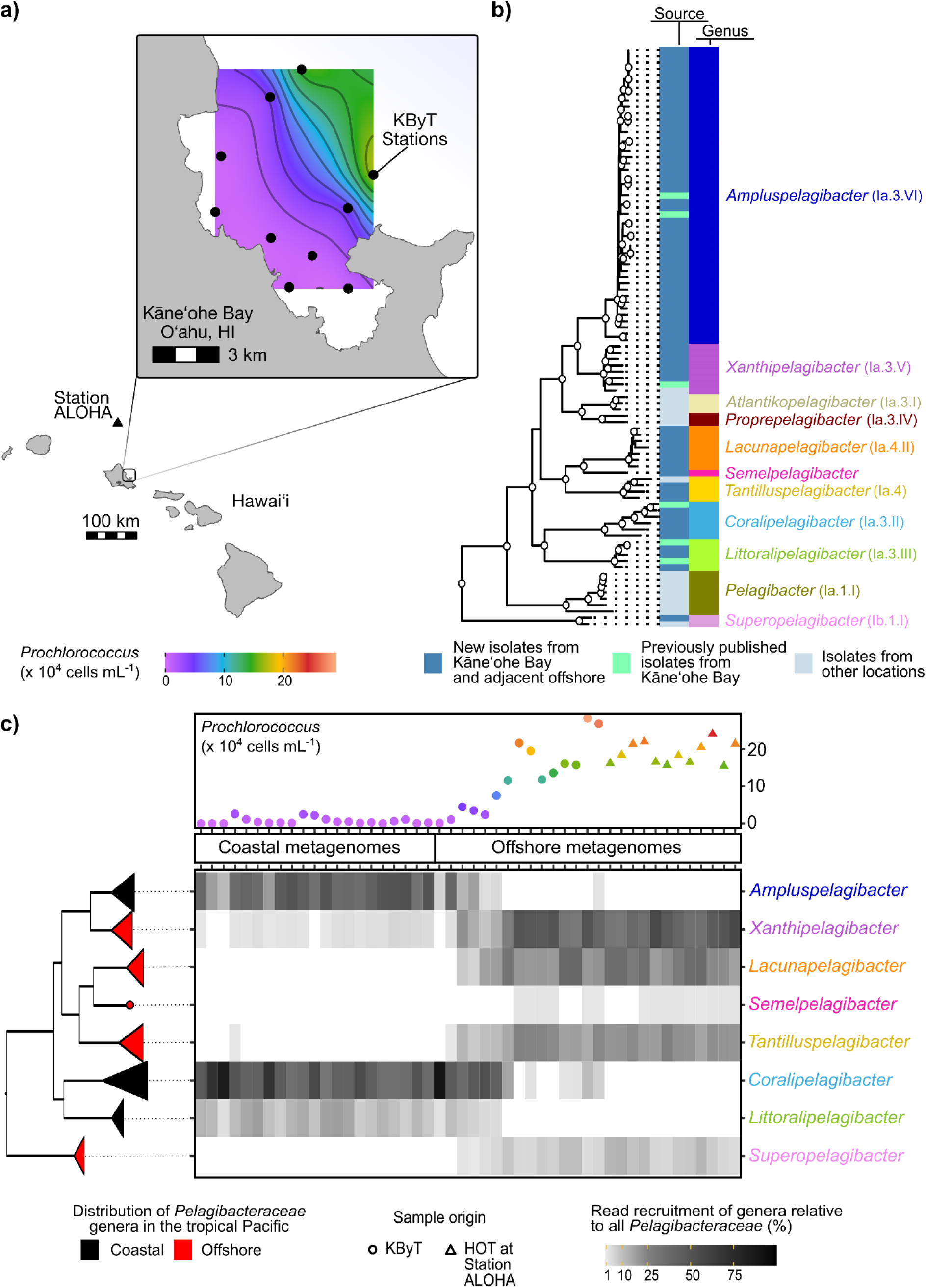
*Pelagibacteraceae* show polyphyletic habitat preferences for coastal or offshore environments across KByT. **a**) The location of the Kāneʻohe Bay Time-series (KByT) sampling stations and the Hawaii Ocean Time-series (HOT) at Station ALOHA. KByT spans a steep biogeochemical gradient, as noted by the dramatic increase in *Prochlorococcus* cellular abundances immediately offshore. **b**) Eleven genera are represented by genome-sequenced *Pelagibacteraceae* isolates, with the majority isolated from coastal Kāneʻohe Bay and the adjacent offshore. Nodes with circles represent ≥90% bootstrap support. Historically referenced clade names for *Pelagibacteraceae* genera are provided in parentheses. **c**) The relative read recruitment of *Pelagibacteraceae* isolate genomes summed at the level of genera for metagenomes from KByT and Station ALOHA. Metagenomes grouped into two clusters by environmental parameters, coastal and offshore, coinciding with the gradient in *Prochlorococcus* cellular abundances. *Pelagibacteraceae* genera showed distinct distribution patterns in the coastal and offshore waters that are distributed polyphyletically. Read recruitment <1% not shown. The order of metagenomes presented follows the order in **Supplementary file 1d**.

We investigated the biogeography of *Pelagibacteraceae* isolate genomes through read recruitment with metagenomes from KByT and Station ALOHA in the adjacent North Pacific Subtropical Gyre. This revealed that the seven genera within the *Pelagibacteraceae* that contained KByT isolate genomes were commonly detected within the KByT system, while the three genera not containing KByT isolates were rarely detected, if at all (**Extended Data Fig. 1b**). Congruent with k-means cluster analysis that grouped the biogeochemical parameters from the metagenomic samples into coastal and offshore clusters (**Extended Data Fig. 3, Supplementary file 1c, Supplementary Information Note 1**), the patterns of genome distribution revealed a clear dichotomy where each genus was either highly present in the offshore or in the coastal environment, but not both (**Fig. 1c, Supplementary file 1d**). Genera with higher abundances in the coastal environment, hereafter referred to as ‘coastal genera’, did not occur within a single monophyletic clade and instead appeared to have evolved multiple times across the evolutionary history of *Pelagibacteraceae* (**Fig. 1c**).

For the most part, the patterns of distribution were consistent for all genomes within the same genus (**Extended Data Fig. 3**), and genomes that belonged to the same genus generally recruited similar proportions of reads from the environment. One exception was the genome HIMB1412 from the genus *Coralipelagibacter* which had the highest relative abundance among all *Pelagibacteraceae* genomes in the coastal environment (19.9±9.4%, mean ± sd, **Extended Data Fig. 3**). While we consistently detected all isolate genomes of *Xanthipelagibacter* in the offshore environment, four of eight genomes (HIMB83, FZCC0015, HIMB1456, HIMB2250) were also frequently detected in the coastal environment (**Extended Data Fig. 3**). Regardless, their relative abundances were low in coastal samples (0.4±0.5%, mean ± sd) and sharply increased offshore (3.5±2.0%, mean ± sd; **Extended Data Fig. 3**), further supporting the characterization of *Xanthipelagibacter* as an offshore genus.

Overall, our metagenomic read recruitment reveals that *Pelagibacteraceae* communities partition between coastal and offshore habitats in the tropical Pacific, that these distinct communities are driven by differences in the distribution of individual *Pelagibacteraceae* genera, and that coastal genera (*Ampluspelagibacter*, *Coralipelagibacter*, *Littoralipelagibacter*) are distributed polyphyletically across the *Pelagibacteraceae*.

### *Pelagibacteraceae* pangenome includes differentially enriched genes and functions within coastal and offshore genera

Next, we sought to characterize the genomic diversity and functional gene content of *Pelagibacteraceae* isolates in order to investigate the potential determinants of fitness that may explain the distinct ecological patterns of distribution. We utilized a pangenomic approach that partitioned all genes across all *Pelagibacteraceae* genomes into *de novo* gene families, or ‘gene clusters’, based on amino acid sequence homology. This analysis resulted in 8,242 gene clusters across all 92 genomes, of which 784 were core among all *Pelagibacteraceae* and made up the majority (52-63%) of gene content found in any individual genome (**Fig. 2a; Supplementary file 2a**). Half of the pangenome was composed of gene clusters that were singletons (4,201 of 8,242; **Fig. 2a**). Between the extremes of gene clusters that occur in every genome and those that occur only in one, we found 206 accessory gene clusters that were ‘genus-specific’ (21±27, mean±sd per genus; **Fig. 2a**; **Supplementary file 2b**) as they were present among all representatives of a single genus, but absent from genomes belonging to any other genus. The 206 genus-specific gene clusters were not distributed uniformly across genera, as 95 were unique to the genus *Superopelagibacter* alone (**Supplementary file 2b**). However, this number is likely an overestimation of the true *Superopelagibacter* genus-specific traits since this particular genus was represented by only two genomes, and the number of genus-specific gene-clusters negatively correlated with the number of genomes in a genus (r^2^adj = 0.37, F=6.29, p = 0.037; Pearson’s correlation). Gene clusters that were core to multiple genera (e.g., present in every genome for two or more genera, but absent in every other) were also not a dominant feature in the pangenome (**Fig. 2a**). They only represented 182 gene clusters, and were generally uniformly distributed across many different groupings of genera (**Fig. 2a**; **Supplementary file 2b**). We could further assign a subset of the gene clusters to coastal or offshore categories based on the environmental preference of the genomes in which they occurred, finding 38 and 176 gene clusters that were specific to coastal genera and offshore genera, respectively. (**Fig. 2a, Supplementary file 2b**).

**Fig 2.**
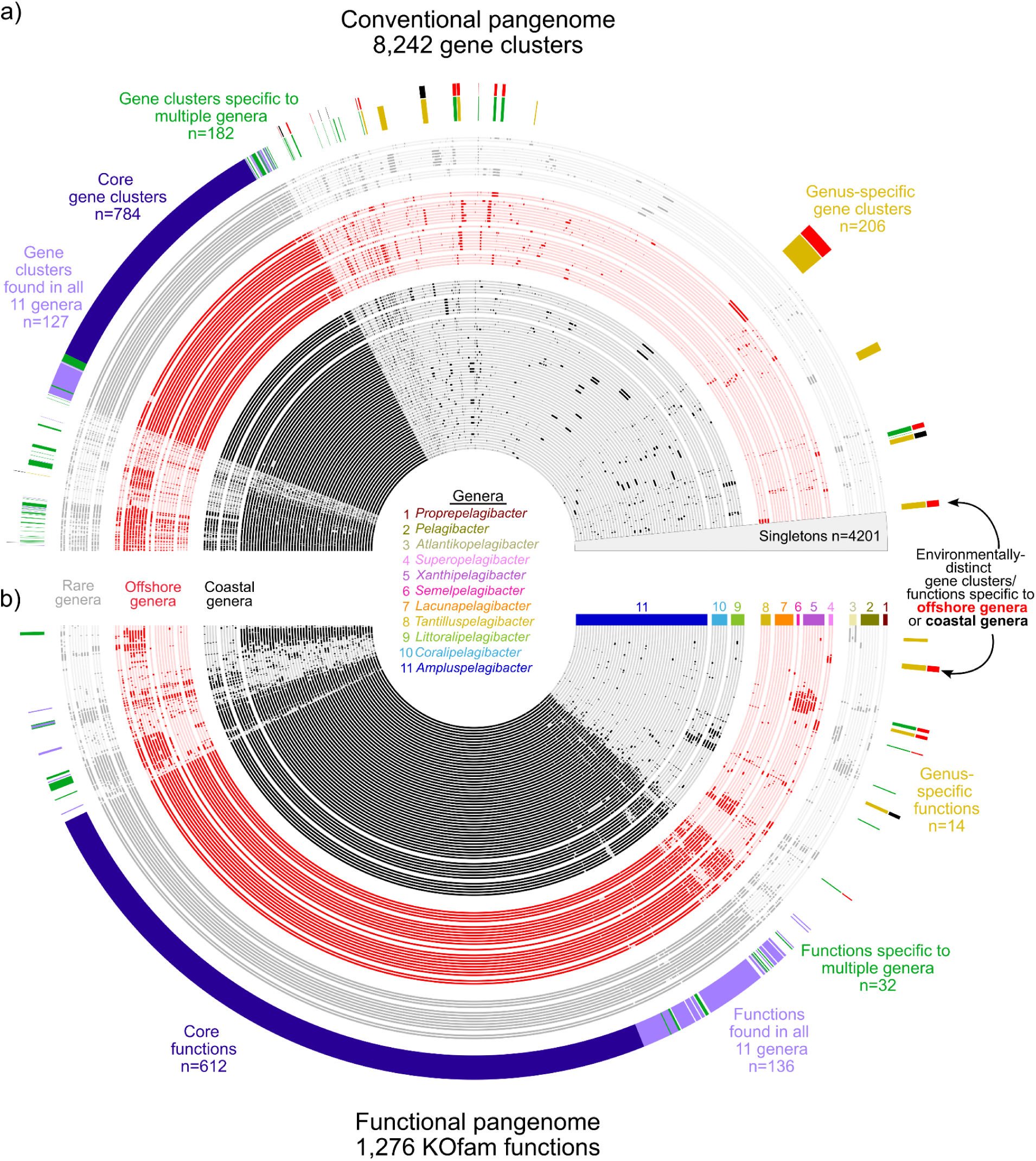
*Pelagibacteraceae* genomes share high overlap in genes and functions among ecologically-distinct groups. The presence and absence of gene clusters and KOfams are shown across **a)** a conventional pangenome and **b)** a functional pagenome, respectively. Genomes (inner rings) are grouped by genus and further grouped by ecological distribution in the tropical Pacific as coastal, offshore, or rare. The distribution of the gene pool and functions in the second most outer layer (e.g., core, genus-specific) show that the gene pool and functions are shared extensively across all 11 genera and 92 genomes. The outer layer shows the sparse environmentally-distinct portions of the gene pool and functional pool that are specific to coastal or offshore genera.

We next focused on genes with well-characterized roles in known metabolisms by examining those that share orthology with genes described in the Kyoto Encyclopedia of Genes and Genomes (KEGG; Kanehisa and Goto 2000) through KOfam models (Aramaki et al. 2020) and generating a ‘functional pangenome’ (**Fig. 2b**). The *Pelagibacteraceae* functional pangenome grouped genes based on functional identity rather than amino acid sequence similarity and consisted of 1,276 KOfam functions, with a large core (612 KOfams; **Fig. 2b**).

Many of the remaining non-core functions were still shared widely across all eleven genera with a relatively uniform distribution (n=136; **Fig. 2b**) or were singletons (n=70). Similar to the conventional pangenome, the functional pangenome contained very few genus-specific functions (14 total among all genera; 0-5 KOfams per genus; **Fig. 2b**) and a limited number of functions that were core to multiple genera (e.g. present in every genome for two or more genera, but absent in every other; 32 KOfams total; **Fig. 2b; Supplementary file 2c**). Only three KOfam functions were specific to coastal genera, while 13 KOfam functions were specific to offshore genera (**Fig. 2a; Supplementary file 2c**).

Pangenomic analysis using conventional and functional approaches revealed a large overlap in gene content among ecologically distinct genera, with subtle but consistent genus-specific features. The high overlap in gene content across the *Pelagibacteraceae* and small number genus-specific gene clusters suggests differentiation between genera of the *Pelagibacteraceae* may be determined by only a small portion of the genome. The polyphyletic distribution of habitat preference (**Fig. 1c**) supports that genes and functions that are linked to success in these different habitats likely evolved multiple times through independent processes. Given this evolutionary trajectory, it is highly plausible that instead of vertical inheritance, horizontal gene transfer led to the acquisition of most of these habitat-specific genes and subsequent high rates of recombination and strong selective forces resulted in the increased prevalence among *Pelagibacteraceae* in a given ecological niche. This process, referred to as gene-specific sweeps, frequently results in a more subtle display of habitat-specific traits (Shapiro et al. 2016), as is observed here. To further examine the potential importance of these environmentally-distinct functions in providing advantages to *Pelagibacteraceae* prevalent in coastal or offshore habitats, we next focused on the subset of the environmentally-distinct functions characterized here that are involved in nutrient resource utilization and cellular stress. Furthermore, in examining the functions involved in nutrient resource utilization and cellular stress, we broadened our investigation beyond the functions that were strictly found in coastal genera or strictly found in offshore genera (e.g. environmentally-distinct functions), to also include functions that were enriched in either coastal or offshore genera.

### Potential determinants of habitat-specificity in *Pelagibacteraceae* include molybdenum utilization and nitrogen and carbon metabolisms

Despite a high amount of water exchange, coastal Kāneʻohe Bay and the adjacent offshore vary in nutrient availability, osmotic conditions (e.g. salinity, temperature, pH), and phytoplankton communities that determine the organic carbon pool (Tucker et al. 2021, 2024), suggesting that this would enforce distinct environmental pressures on coastal and offshore *Pelagibacteraceae* genera. To characterize ecologically-relevant genes that may support coastal and offshore habitat-preferences of *Pelagibacteraceae* across the KByT system, we used an enrichment analysis to examine metabolic traits related to nutrient acquisition and cellular stress that were differentially distributed across genomes from either coastal or offshore genera. Genes that were core across genera of the *Pelagibacteraceae* or did not show easily distinguishable distributions (e.g. driven by phylogeny or environment) are further discussed in **Supplementary Information Note 2**. Unless noted otherwise, the following metabolic traits were found outside of hyper-variable regions of genomes and thus represent genes that are encoded within the relatively stable genomic backbones of a given genus or set of genera.

### Molybdenum enzyme utilization

Molybdenum enzymes are nearly ubiquitous among organisms from all domains of life, catalyze important oxidation-reduction reactions involved in carbon, sulfur, and nitrogen metabolisms, and require a cofactor scaffold to hold the molybdenum in place (Zhang and Gladyshev 2008). While most *Pelagibacteraceae* genomes contained genes involved in molybdenum cofactor biosynthesis as well as molybdenum cofactor-dependent enzymes (**Fig. 3a, Supplementary file 2d**), these genes were missing from two offshore genera, *Lacunapelagibacter* and *Tantilluspelagibacter*. In contrast to the lack of molybdenum enzymes found in the offshore *Lacunapelagibacter* and *Tantilluspelagibacter*, other offshore genera, *Xanthipelagibacter* and *Semelpelagibacter*, appeared to increase reliance on molybdenum-dependent enzymes by uniquely harboring genes encoding molybdenum-dependent enzymes associated with the oxidation of purines (xanthine; *xdhAB*) (**Fig. 3a, Supplementary file 2d, Supplementary Note 3**). The distribution of molybdenum enzymes are polyphyletically distributed among offshore genera, where some offshore groups possess no molybdenum enzymes while others harbor all the molybdenum enzymes detected within *Pelagibacteraceae* (**Fig. 3a**). We hypothesize that the absence of molybdenum enzymes within the offshore *Lacunapelagibacter* and *Tantilluspelagibacter*, which have some of the smallest genomes within *Pelagibacteraceae* (∼1.2Mbp; **Supplemental file 1a**), is likely driven by streamlining selection to minimize metabolic requirements and genome size (Giovannoni et al. 2005, 2014).

**Fig. 3.**
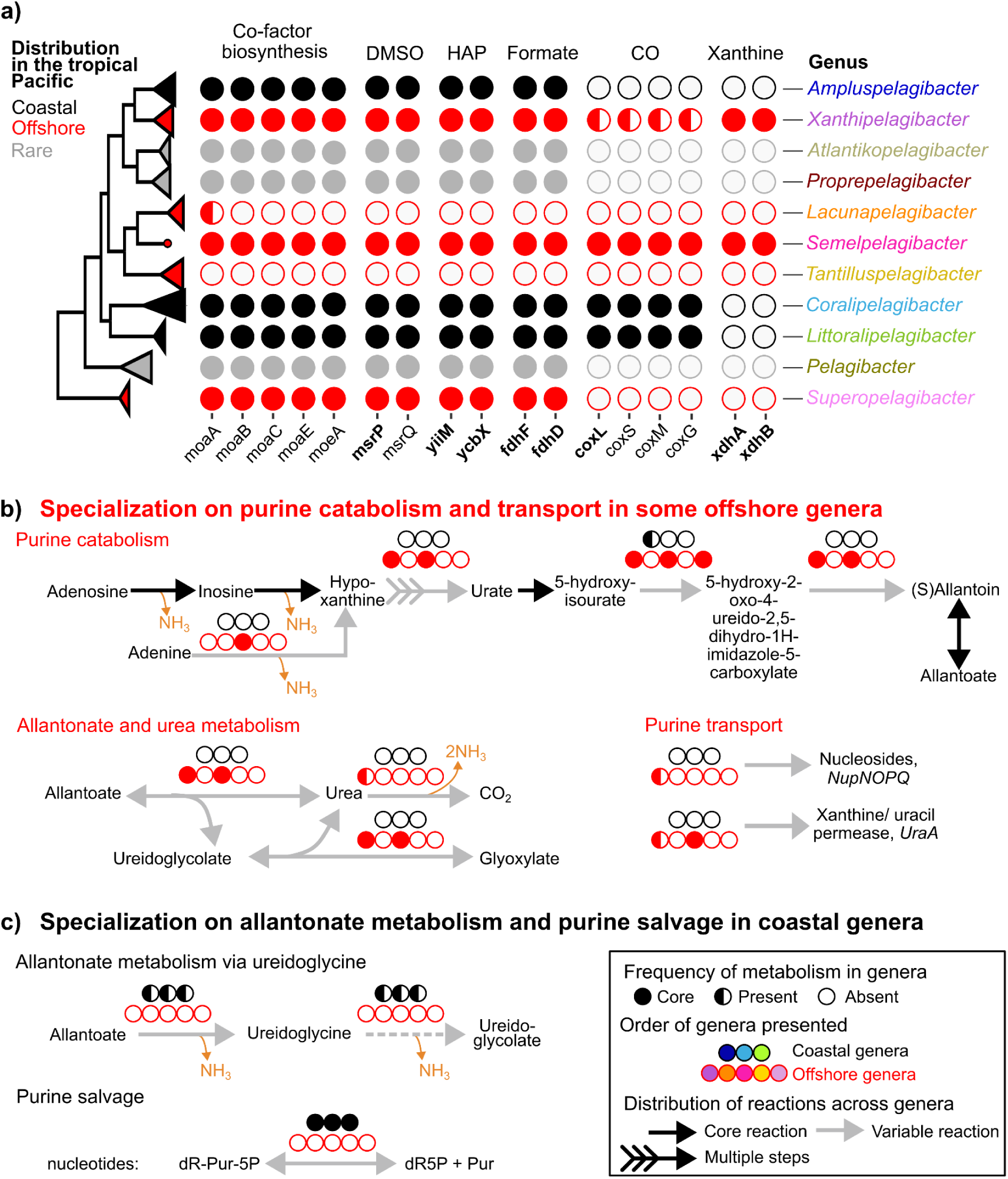
*Pelagibacteraceae* genera fill distinct niches in molybdenum enzyme utilization. **a)** The distribution of molybdenum cofactor (MoCo) biosynthesis genes and genes involved in metabolisms that require a molybdenum cofactor across *Pelagibacteraceae* genera. Metabolisms within *Pelagibacteraceae* that require a molybdenum cofactor included those involved in the repair of methionine sulfoxides and the catalytic subunit of periplasmic dimethylsulfoxide (DMSO; *msrPQ*), the detoxification of 6-N-hydroxylaminopurine (HAP) to adenine (*yiiM*, *ycbX*), the oxidation of formate to carbon dioxide (*fdhFD*), the oxidation of carbon monoxide (*coxSMLG*), and the oxidation of purines via xanthine dehydrogenases (*xdhAB*). Genes that are predicted to represent proteins with molybdenum cofactor binding sites are noted in bold. **b)** Genes encoding for purine catabolism and transport are only present in some offshore genera. **c)** Genes encoding for allantonate metabolism and purine salvage are present only in coastal genera. MoCo: molybdenum cofactor, DMSO: dimethylsulfoxide, HAP: hydroxylaminopurine, CO: carbon monoxide, NH_3_: ammonia, dR-Pur-5P: purine 2’-deoxyribonucleoside 5’-monophosphate, Pur: purine base, dRib5P: 2’-deoxyribonucleoside 5’-monophosphate, (d)R-Pur-1P: beta-(deoxy)ribonucleosides.

### Nitrogen metabolisms

Purines are abundant in aquatic environments (Berman and Bronk 2003), contain a high number of nitrogen atoms per molecule (Kornberg 1974), and likely contribute to niche-differentiation between coastal and offshore SAR11 (Braakman et al. 2025). Our metabolic reconstructions show that offshore genera *Xanthipelagibacter* and *Semelpelagibacter* harbored genes to catabolize purines through the utilization of a molybdenum cofactor-dependent xanthine dehydrogenase (**Fig. 3b, Supplementary file 2e**), as well as genes with functions predicted to support the efficient transport of nucleotides (*Xanthipelagibacter*: *nupNOPQ* system; *Xanthipelagibacter* and *Semelpelagibacter*: xanthine/uracil permease *uraA*). To make hypoxanthine available for degradation, *Semelpelagibacter* can convert adenine to hypoxanthine using adenine deaminase (*ade*) and both genera *Semelpelagibacter* and *Xanthipelagibacter* also uniquely possess a tRNA(adenine34) deaminase (*tadA*), which can convert adenine to hypoxanthine (**Fig. 3b**). Most *Xanthipelagibacter* genomes (6 of 8) also harbor a urease system (*ureABCDEFG*) to liberate ammonia in the final steps of the xanthine degradation pathway.

In contrast to the purine degradation pathways found within the offshore genera, we found coastal genera specialize on the degradation of allatonate, a bi-product of purine metabolism, as well as the recycling of nucleotides. Members of the coastal genus *Ampluspelagibacter* (and a single *Coralipelagibacter* genome, HIMB5) have the functional capacity to degrade allantoate to (S)-ureiodoglycine and ammonia via an allantoate deiminase and ureidoglycine to ureidoglycolate and ammonia, via a nucleophile (Ntn)-hydrolase (**Fig. 3c, Supplementary file 2e**). It is important to note that the allantoin degradation genes found in coastal genomes characterized here appear to provide a source of nitrogen, and differ from those found in the offshore genera. All coastal genera possess a deoxynucleoside 5-monophosphate N-glycosidase (*rcl*) that putatively breaks the N-glycosidic bond of purine nucleotides (i.e. purine 2’-deoxyribonucleoside 5’-monophosphates; **Fig. 3c**). This differs from the purine nucleoside salvage pathway core to both coastal and offshore genera, which requires orthophosphate to cleave the N-glycosidic bond of beta-(deoxy)ribonucleosides to yield alpha-(deoxy)ribose 1-phosphate (**Supplementary file 2e**). All *Pelagibacteraceae* are missing genes to utilize deoxyribose-5-phosphate or deoxyribose-1-phosphate via a deoxyribose-phosphate aldolase (*deoC*). However, ribose-1-phosphate could be catabolized to D-fructose-6P and D-glyceraldehyde-3P (**Supplementary file 2e**).

In addition to differences in purine metabolism, offshore genera uniquely harbor a transaminase (*agxt*) that may help these cells to overcome auxotrophies for the biosynthesis of the amino acids glycine and serine (**Fig. 4, Supplementary file 2f**), which are common features across *Pelagibacteraceae*, but likely restrict important biosynthetic pathways including protein synthesis and central carbon metabolism (Tripp et al. 2009). Offshore genera also uniquely contain genes involved in glutamate deamination, glutathione and carnitine metabolism, and the synthesis of glucosylglycerate, which is used as an alternative osmolyte under nitrogen-limited conditions in cyanobacteria (Klähn et al. 2010). In contrast, the coastal genera contain nitrogen metabolisms generally involved in the deamination of various amino acids (e.g. hydroxyprolines, proline, ornithine, and threonine). A subset of coastal *Ampluspelagibacter* genomes also have the potential to metabolize the polyamines putrescine and spermidine via *puuBCD* (**Fig. 4**). The puuD gene was located in hypervariable regions within coastal *Ampluspelagibacter* genomes.

**Fig. 4.**
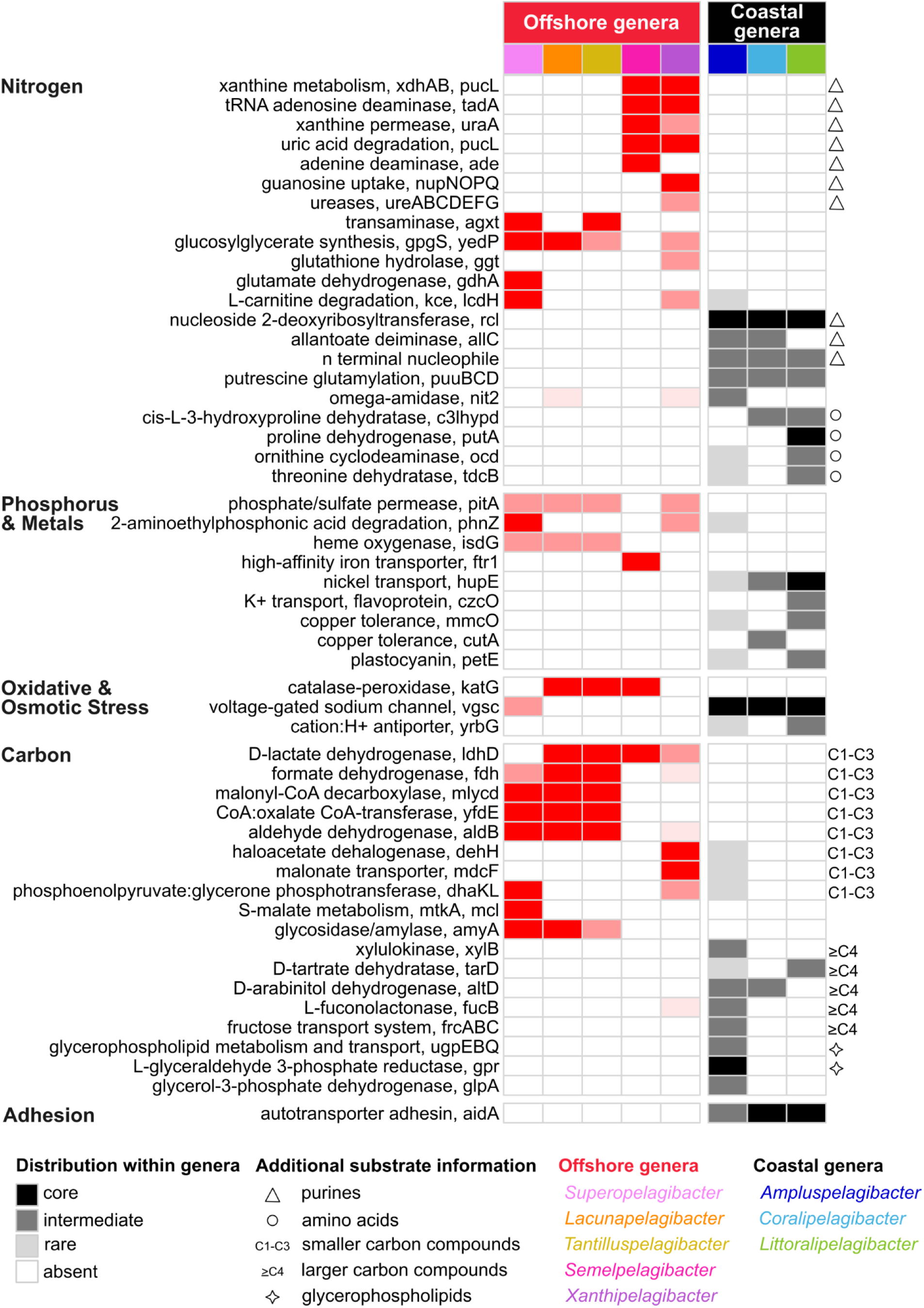
Metabolic determinants of habitat-specificity in *Pelagibacteraceae*. The distribution of metabolic genes associated with nutrient acquisition of carbon, nitrogen, phosphorus, and metals and oxidative and osmotic cellular stress vary among coastal and offshore *Pelagibacteraceae* genera. Each column represents a genus, organized by genera that have coastal (black) and offshore (red) biogeographical distributions. Each row represents a metabolic gene(s) organized by their role in carbon, nitrogen, phosphorus, or metal acquisition or oxidative and osmotic cellular stress. The intensity of color indicates the prevelance of the metabolic gene(s) in each genus.

### Phosphate & Metals

Some offshore *Pelagibacteraceae* genera exhibit a reduced genomic capacity to transport phosphate relative to coastal genera. High-affinity phosphate acquisition genes (*pstSCAB-phoU*; *phoB*; *phoR*) were core among coastal genera but, while present in some genomes, were not core among the offshore genera. Offshore genomes lacking this system appear to rely solely on the *pitA* gene, a presumably lower-affinity proton-motive force permease of phosphate or sulfur (**Fig. 4, Supplementary file 2f**). Some offshore genera contain an increased genomic capacity to transport or capture iron relative to coastal genera. While all *Pelagibacteraceae* contain an iron (III) transport system (*afuAB*), the offshore *Semelpelagibacter* genus alone possesses a high-affinity iron permease (*ftr1*) (**Fig. 4, Supplementary file 2f)**. Offshore genera *Superopelagibacter*, *Lacunapelagibacter*, and *Tantilluspelagibacter* also contain a heme oxidase, which could catalyze the oxidative degradation of the heme porphyrin ring to release iron.

Unique genes to utilize copper, potassium, and nickel were found among the coastal genera (**Fig. 4)**. We observed a copper-containing plastocyanin gene within a small subset of coastal genomes (**Fig. 4; Supplementary file 2f**). Plastocyanins serve as an electron carrier from cytochrome *f* to photosystem I and are generally found within eukaryotic phytoplankton and cyanobacteria (Castell et al. 2021). Future studies are needed to understand the functional role of plastocyanins in *Pelagibacteraceae*, although one potential use could be the storage of copper to cope with periods of reduced availability (Peers and Price 2006).

### Cellular stress

Differences in the capacities to respond to osmotic and oxidative stresses were found among offshore and coastal *Pelagibacteraceae* genera. Most offshore genera shared a catalase-peroxidase (*katG*) that may help to cope with oxidative stress and has also been suggested to play a role in the co-evolution of marine microbial communities (Morris et al. 2012) (**Fig. 4, Supplementary file 2f**). Coastal genera contained two genes involved in osmotic stress that were not prevalent among offshore genomes: a voltage-gated sodium channel (*vgsc*) and a cation:H+antiporter (*yrbG*) (**Fig. 4, Supplementary file 2f**). Voltage-gated sodium channels utilize a structurally complex gating system to physically open and close the pore in response to changes in fluid shear or membrane stretch from dehydration (Ren et al. 2001; Strege et al. 2023), and may be particularly useful in coastal Kāneʻohe Bay where spatiotemporal shifts in salinity are common (Yeo et al. 2013).

### Carbon metabolisms

Carbon metabolisms unique to coastal and offshore genera were remarkably distinct. SAR11’s specialization on low-molecular weight, labile carbon sources has likely facilitated the success of this marine oligotroph in open oceans where carbon substrates are limited in availability and under high competition (Sun et al. 2011; Giovannoni 2017; Moore et al. 2022). Consistent with these observations, the carbon metabolisms shared by offshore genera did not rely on glycolysis and involved low-molecular-weight compounds (one-carbon to three-carbon; C1-C3) that are common molecules and/or typical waste products in the marine environment.

This includes the genetic potential to metabolize organic acids such as D-lactate, oxalate, haloacetate, and formate, as well as diverse aldehydes (**Fig. 4**, **Supplementary file 2f**). In contrast, coastal genera contain gene content to specialize in the degradation of relatively larger carbon compounds (C4-C6), including the sugars D-xylulose and D-ribulose via *xylB* and sugar alcohols arabinitol and mannitol via an arabinitol dehydrogenase, both of which subsequently enter glycolysis through the non-phosphorylative Entner–Doudoroff (np-ED) pathway. Some coastal groups also have the capacity to metabolize the sugar acids D-tartrate and malate via *tarD* (**Fig. 4; Supplementary file 2f**). Genomes belonging to the coastal genus *Ampluspelagibacter*, which have the largest genomes among known *Pelagibacteraceae*, harbored gene content potentially involved in the transport and metabolism of glycerophospholipids and sn-glycerol-3-phosphate (*upgEBQ*, *gpr*, *glpA*), the capacity to metabolize L-fucose, D-arabinose, and L-xylose to pyruvate through a L-fucono-1,5-lactonase (*fucB*) and the glycolytic Entner-Doudoroff (ED) glycolytic pathway, and a fructose transport system (*frcABC*) (**Fig.4; Supplementary file 2f**). The ED glycolytic pathway was core among *Ampluspelagibacter*, *Xanthipelagibacter*, and *Proprepelagibacter* genomes and variable across genomes in other genera (**Supplementary file 2f**). The increased prevalence of sugar metabolisms among coastal genera, especially the *Ampluspelagibacter* genus, suggests broadened metabolic versatility among coastal *Pelagibacteraceae* in response to a diverse and/or more readily available set of organic carbons found within coastal Kāneʻohe Bay relative to offshore waters. The distinct distributions of carbon metabolisms resolved here also suggests fine-scale niche partitioning along the sugar-acid spectrum within *Pelagibacteraceae* (Gralka et al. 2023).

### Adhesion

Finally, while investigating the metabolic differences between coastal and offshore genera, we observed homologs of a large gene (∼5,000 to 9,500 bp) annotated as an autotransporter adhesin that was almost always present among coastal genomes and absent in all offshore genomes (**Fig. 4**). Autotransporters provide a simple and relatively minimal mechanism for the delivery of a passenger protein to the surface of Gram-negative bacteria (Leyton et al. 2012), and can vary greatly in size due to changes in the number of repeating sequences in the passenger protein (Doyle et al. 2015). Because repeating regions cause difficulty for short-read assemblers such as those used in our genome assemblies (Freel et al. 2024), we further established confidence in the assembly of this gene in our coastal genomes by successfully finding homologs in long-read sequencing libraries from the coastal Kāneʻohe Bay environment and comparing protein structure models of genes annotated from the long-read data to a known autotransporter (**Extended Data Fig. 4; Supplementary Information Note 4**). We also examined the genome context of the gene, which was always positioned next to genes involved in type IV pilus systems and/or type II secretory systems and sometimes, but not always, within hyper-variable regions (**Extended Data Fig. 4, Supplementary Information Note 4, Supplementary file 2f**).

Adhesion to surfaces has not been observed in *Pelagibacteraceae* in culture or through environmental studies, although long filaments believed to be pili were observed within dividing cells of *Pelagibacteraceae* (Zhao et al. 2017) and are associated with multiple processes, including adhesion to surfaces, transformation competence, and DNA uptake (Hazes and Frost 2008; Ellison et al. 2018). While speculative, it is possible that adhesion by the autotransporter adhesin gene may be involved in the uptake of nutrients also in *Pelagibacteraceae* (Meuskens et al. 2019). The coastal distribution of the autotransporter gene suggests that the advantage of having this large of a gene is worth the trade-off of maintenance in more nutrient-rich coastal environments, but potentially not in nutrient-limited offshore environments.

In summary, our investigation of the distribution of genes related to nutrient acquisition and cellular stress that are unique to coastal and offshore genera reveals numerous subtle differences in metabolic traits driven by the presence or absence of single genes (e.g. high affinity iron transporters, voltage-gated sodium channel genes), but few differences in multi-gene metabolic pathways (e.g. purine metabolism, molybdenum cofactor biosynthesis, fucose metabolism). Differences in genes related to nutrient acquisition were predominantly related to organic carbon and nitrogen metabolisms, and help to further define the unique and diverse roles *Pelagibacteraceae* play in oceanic biogeochemical cycles. The distribution of these metabolic traits generally do not follow phylogenetic distributions, but environmental patterns, suggesting that these metabolisms may be driven by selective pressures from the environment.

### Gene determinants of habitat-specificity are under relatively higher selective pressures compared to non-diagnostic genes

Our analyses indicated a relatively small number of functions in the *Pelagibacteraceae* appear to be associated with its strict niche partitioning. Assuming that the requirement to perform optimally may be higher for genes that determine fitness to particular lifestyles, we hypothesized that metabolic traits that have distinct environmental distributions should be under higher selective pressures compared to other genes in the environment. To test this hypothesis we examined the proportion of non-synonymous to synonymous (pN/pS) sites per gene across *Pelagibacteraceae* genomes using four deeply sequenced metagenomic samples from the KByT system.

To define the set of genomes to be included in this analysis, we first evaluated whether the gene-level selective pressures were uniform across genomes from the same genus. We examined variation in pN/pS across genes with shared KOfam assignments for genomes belonging to the genus *Coralipelagibacter,* which has a low minimum gANI (84.5%, n=6) and is abundant in the tropical Pacific. Gene-to-gene variation explained 82% of the variation, while sample-to-sample variation and genome-to-genome variation only explained 0.26 and 0.09%, respectively (ANOVA, **Supplemental file 3a**). The little genome-to-genome variation showed that genomes within the same genus likely experience similar patterns of selection on shared genes, and supported using a single genome as a representative per genus. Thus, our downstream analyses utilized a single genome for each genus in *Pelagibacteraceae*, except *Atlantikopelagibacter*, *Pelagibacter*, *Proprepelagibacter*, and *Semelpelagibacter*, which lacked sufficient coverage in the KByT system (**Extended Data Fig. 1, Supplemental file 3b**).

Across the *Pelagibacteraceae* genomes examined, strong purifying selection (pN/pS ≪ 1) was a general characteristic of most genes as observed previously by Kiefl et al., (2023), where over 75% of genes in each genome representative had a pN/pS value of ≤ 0.1 (**Supplemental file 3c**). However, not all genes experienced similar magnitudes of purifying selection, with gene-to-gene variation in pN/pS explaining between 88-99% of the variation in pN/pS across genomes (n=7, ANOVA, **Supplemental file 3d**). Sample-to-sample variation in pN/pS values varied for most genomes, however it explained a very small proportion of the overall variation (<0.5%, n=7, ANOVA). Gene coverage showed no correlation between per gene values of pN/pS (absolute value of r=0.02-0.28; Pearson correlation coefficient; **Extended Data Fig. 5**). Gene coverage explained an extremely small portion of the overall pN/pS variation (<0.07%, n=7, ANOVA, **Supplemental file 3d**), which indicates that pN/pS values are unlikely to be driven by artifacts associated with variation in coverage.

We next evaluated the selective pressures on genes that were differentially distributed across coastal and offshore *Pelagibacteraceae* genera (**Fig. 5; Extended Data Fig. 6)**. The differentially distributed genes with the highest selective pressures in offshore genera included a malonyl-CoA decarboxylase (*mlycd;* pN/pS= 0.019±0.010, mean ± sd), an aldehyde dehydrogenase (*aldB*; pN/pS= 0.033±0.018), a formate dehydrogenase (*fdh;*pN/pS=0.0127±0.001), a D-lactate dehydrogenase (*ldhD*; pN/pS=0.018±0.003) and an alanine-glyoxylate / serine-glyoxylate / serine-pyruvate transaminase (*agxt*; pN/pS= 0.022±0.008) (**Fig. 5**). These genes are generally involved in the metabolism of small and common carbon substrates and the capacity to produce glycine and serine, which are auxotrophies in other *Pelagibacteraceae*. Within *Xanthipelagibacter*, genes involved in purine uptake (*uraA*, pN/pS=0.040±0.001; *nupNOPQ*, pN/pS=0.045±0.010) experienced higher purifying selection than purine degradation pathways (*xdhAB*, pN/pS=0.084±0.028), which is in line with previous observations that high-affinity transport is key to *Pelagibacteraceae*’s success as an oligotroph (Noell and Giovannoni 2019; Clifton et al. 2024). The low pN/pS values for these distinct carbon and nitrogen related genes support the hypothesis that these metabolisms provide a fitness advantage in offshore waters.

**Fig. 5.**
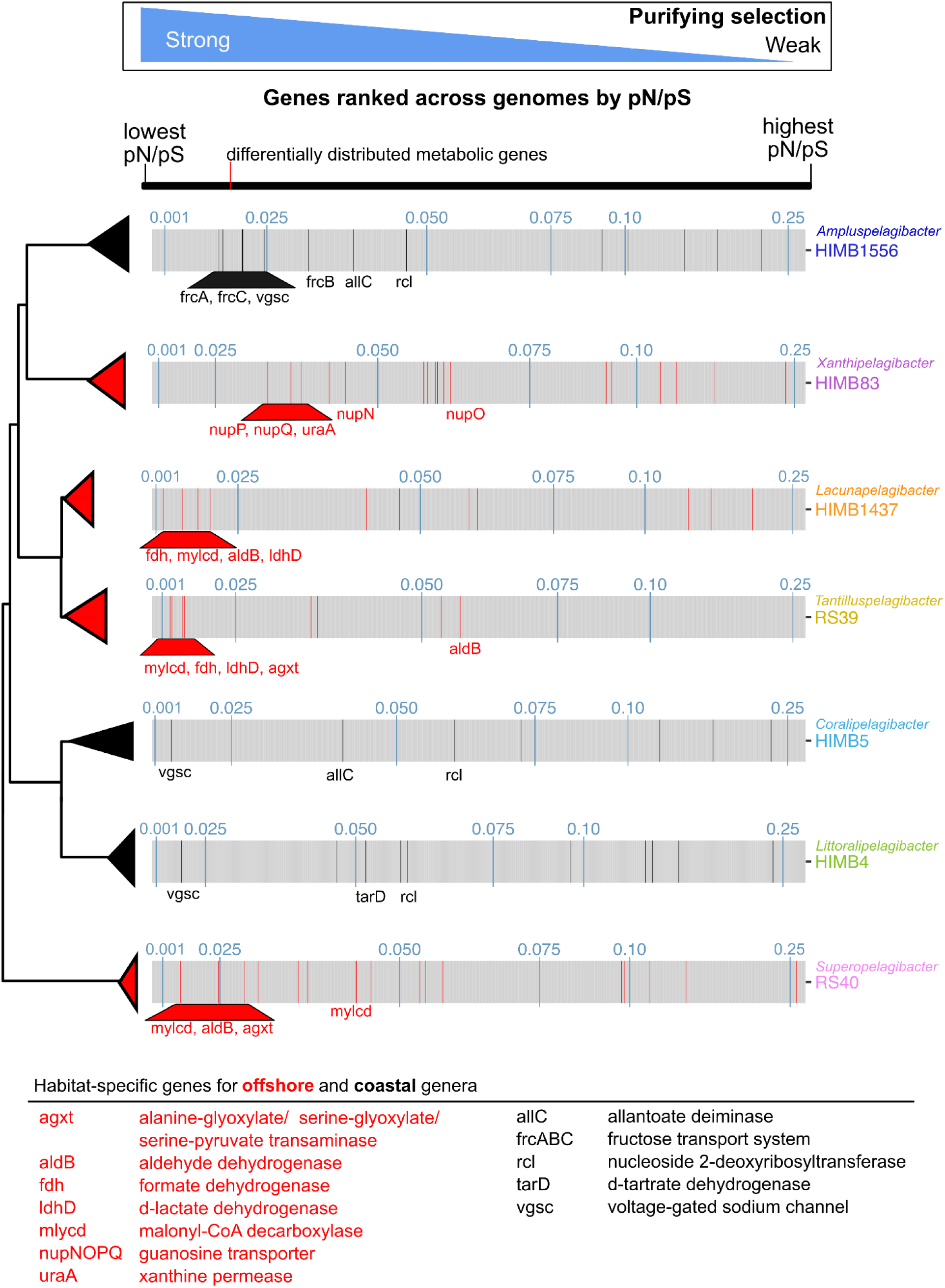
Habitat-specific genes are under relatively higher selective pressures compared to non-diagnostic genes. Across each genome, pN/pS values per gene are ranked from lowest pN/pS value (high purifying selection) to highest pN/pS value (low purifying selection). Genes that were characterized as differentially distributed between offshore and coastal *Pelagibacteraceae* genera in Fig. 4 are colored red (offshore) or black (coastal) across the genomes. The gene names are noted and the description of the gene provided in the key for genes that are discussed in the text. The full suite of gene names and descriptions can be found in **Extended Data Fig. 6.** pN/pS values of 0.01, 0.025, 0.05, 0.075, 0.1, and 0.25 are shown in blue and genomes are ordered by their phylogenomic relationships. pN/pS: proportion of non-synonymous to synonymous.

Among the genes unique to coastal genera, the voltage-gated sodium channel was under high selective pressure (*vgsc*, pN/pS=0.016±0.004). Coastal Kāneʻohe Bay harbors a larger range of salinity, temperature, and pH conditions compared to the adjacent offshore (Yeo et al. 2013; Tucker et al. 2021, 2024), and thus the voltage-gated sodium channels may be critical for coastal genera as they rapidly regulate ionic composition (Ren et al. 2001). The allantoin deiminase (*allC*, pN/pS=0.039±0.002) and nucleoside 2-deoxyribosyltransferase (*rcl*, pN/pS=0.054±0.006) were also under moderately high selective pressures in coastal genomes, further supporting the specialization of coastal genera on allantonate and nucleotides and a separation in purine metabolism niche from offshore genera. Three sugar transporter genes (*frcABC*, pN/pS=0.024±0.006) within *Ampluspelagibacter* and a tartrate dehydratase (*tarD*, pN/pS=0.051±0.001) within *Littoralipelagibacter* exhibited low pN/pS values suggesting that sugar metabolisms and transporters are not only a more common genomic feature in the coastal genera, but also likely essential for their fitness.

Finally, to understand which *Pelagibacteraceae* core genes were under the highest selective pressures in the KByT system, we examined the mean pN/pS values for core genes found across the same seven genomes. Not surprisingly, ribosomal proteins were found among the genes with the highest purifying selection (low pN/pS values; **Table 2**). Genes related to baseline functions of the cell including general energy production and conversation, translational, ribosomal structure and biogenesis, transcription, and nucleotide transport and metabolism were also under high selective pressure (**Table 2**). Two genes involved in regulatory systems for responses to acidity and glycine were among the top 25 core genes with the most purifying selection. The low pN/pS values for these genes supports that although *Pelagibacteraceae* may have few regulatory genes generally (Giovannoni et al. 2014), the ones it does have likely impact the performance and survival of *Pelagibacteraceae* cells.

**Table 2.** Core *Pelagibacteraceae* genes under high selective pressures in the KByT system. . The lowest 25 mean pN/pS values for core *Pelagibacteraceae* genes with shared KOfam annotations examined from genomes representing the seven *Pelagibacteraceae* genera that are highly abundant in the tropical Pacific. pN/pS: proportion of non-synonymous to synonymous. Enzymes involved in regulatory processes are denoted by a pound sign (#).

| Accession | Description | pN/pS<br>(mean±s.d.) |
| --- | --- | --- |
| K02946 | small subunit ribosomal protein S10 | 0.012±0.004 |
| K00340 | NADH-quinone oxidoreductase subunit K | 0.013±0.008 |
| K03560 | biopolymer transport protein TolR | 0.014±0.007 |
| K03628 | transcription termination factor Rho | 0.014±0.004 |
| K02078 | acyl carrier protein | 0.014±0.007 |
| K03544 | ATP-dependent Clp protease, subunit ClpX | 0.017±0.004 |
| K00053 | ketol-acid reductoisomerase | 0.017±0.005 |
| K00395 | adenylylsulfate reductase, subunit B | 0.018±0.008 |
| K00394 | adenylylsulfate reductase, subunit A | 0.018±0.005 |
| K00412 | ubiquinol-cytochrome c reductase cytochrome b subunit | 0.018±0.006 |
| K02274 | cytochrome c oxidase subunit I | 0.018±0.005 |
| K02982 | small subunit ribosomal protein S3 | 0.019±0.007 |
| K03566# | LysR family transcriptional regulator, glycine cleavage system<br>transcriptional activator | 0.019±0.011 |
| K01494 | dCTP deaminase | 0.019±0.006 |
| K02878 | large subunit ribosomal protein L16 | 0.019±0.008 |
| K03798 | cell division protease FtsH | 0.019±0.005 |
| K03702 | excinuclease ABC subunit B | 0.020±0.008 |
| K00330 | NADH-quinone oxidoreductase subunit A | 0.020±0.010 |
| K03569 | rod shape-determining protein MreB | 0.020±0.011 |
| K02874 | large subunit ribosomal protein L14 | 0.020±0.011 |
| K14981# | two-component system, OmpR family, response regulator ChvI | 0.020±0.01 |
| K02988 | small subunit ribosomal protein S5 | 0.021±0.005 |
| K03046 | DNA-directed RNA polymerase subunit beta | 0.021±0.003 |
| K03701 | excinuclease ABC subunit A | 0.021±0.003 |
| K00331 | NADH-quinone oxidoreductase subunit B | 0.021±0.006 |

Only two genes related to nutrient acquisition were found among the top 25 core genes with the highest purifying selection. This included both subunits of an adenylylsulfate reductase (*aprAB*), which is involved in dissimilatory sulfur metabolism through the introconversion of adenylyl sulfate (APS) to sulfite. Incomplete dissimilatory and assimilatory sulfate reduction pathways were core among all *Pelagibacteraceae* examined in this study (**Extended Data Fig. 7**), in line with past analysis of *Pelagibacteraceae* genomes (Tripp et al. 2008). As reported by Smith et al. (2016), adenylylsulfate reductase likely removes sulfite that accumulates during organic sulfur compound degradation. Further supporting this hypothesis, we found multiple core or near core genes involved in the production of sulfite from reduced organosulfur compounds: (2R)-sulfolactate sulfo-lyase (*suyAB*), a sulfoacetaldehyde acetyltransferase (*xsc*), and a sulfur dioxygenase (**Extended Data Fig. 7**). Our analyses reveal the genetic potential to metabolize a wide diversity of reduced organosulfur compounds as alternative sources of sulfur and that adenylylsulfate reductases (*aprAB*) likely plays an important role in accommodating *Pelagibacteraceae*’s unique sulfur metabolisms.

## Discussion

Through genomic and metagenomic characterizations of a recently expanded collection of SAR11 isolates (Freel et al. 2024), our study reveals a polyphyletic distribution of coastal ocean versus offshore habitat specialists among closely related SAR11 populations, where the emergence of key genetic features that underlie this strong habitat preferences seem to have occurred through independent evolutionary events rather than transfer from a single common ancestor. Genera that share the same environment (e.g. coastal or offshore) have also accumulated shared and unique gene content that is predominantly involved in the metabolism of organic carbon and the acquisition of nitrogen from organic sources. A subset of these metabolic genes were under high purifying selection, emphasizing the importance of these functions to the fitness of distinct *Pelagibacteraceae* genera and highlighting potential determinants of niche differentiation of *Pelagibacteraceae* in coastal and offshore environments.

The cohesion of both ecological and genetic diversity observed within genera of *Pelagibacteraceae* most closely resembles ecotypes: ecologically homogeneous groups of closely related bacteria whose genetic diversity is guided by cohesive forces of selection, recombination, and genetic drift (Cohan 2006; Koeppel et al. 2008). Ecotypic differentiation has previously been suggested to describe the spatial and temporal variation partitioning genetic and genomic SAR11 diversity (Field et al. 1997; Schwalbach et al. 2010; Brown et al. 2012; Vergin et al. 2013; Tsementzi et al. 2019; Haro-Moreno et al. 2020; Kraemer et al. 2020; Tucker et al. 2021; Larkin et al. 2023; Freel et al. 2024), however the underlying evolutionary processes or functional differences contributing to this diversification have rarely been linked to patterns of distribution. Our findings reveal evolutionary drivers behind ecotypic differentiation by explaining distinct ecological distribution patterns with differences in gene content, metabolic potential, and selective pressures across ecotypes. The use of a constrained model system that provided both metagenomic sequence data as well as isolate genomes from local *Pelagibacteraceae* populations was paramount to being able to resolve signatures of environmental selection in sympatric and parapatric SAR11 populations. The power of this approach is further demonstrated in the fact that most of the *Pelagibacteraceae* ecotypes defined in this constrained system as having preferences for coastal oceans or the open ocean appear to hold when global read recruitment data were examined (Freel et al. 2024).

Our results suggest that *Pelagibacteraceae* has transitioned between offshore and coastal environments multiple times, with a handful of genes that support these lifestyles likely acquired through multiple independent processes. The small number of adaptive genes are shared among ecologically similar, but polyphyletically-distributed genera, where genomes have maintained a substantial amount of genetic diversity. Given the distribution patterns of the adaptive genes and the genetic diversity maintained within the populations that carry them, the proliferation of adaptive genes unlikely occurred via genome-wide selective sweep (Shapiro et al. 2016), a process that describes the clonal expansion of an individual subpopulation that carries an adaptive gene leading to a genome-wide reduction in genetic diversity. Instead, the maintenance of genetic diversity and the relatively strong purifying selection to maintain the functional identity of these adaptive genes is consistent with a gene-specific selective sweep (Falush et al. 2001; Whitaker et al. 2005; Shapiro et al. 2012; Rosen et al. 2015; Bendall et al. 2016).

Consistent with previous studies that have identified a handful of genes responsible for habitat-specific adaptation (Shapiro et al. 2012; Bendall et al. 2016; Delmont and Eren 2018), the gene content differences attributed to stable ecological speciation here are generally found within the relatively stable genomic backbone, rather than within hyper-variable genomic islands. This is a critical observation as it links eco-evolutionary processes to specific portions of the immense genomic diversity that exists within *Pelagibacteraceae*, and underscores the importance of analyses informed by genomic architecture. Future studies that examine the contribution of gene content found within hyper-variable islands to stable ecological speciation as opposed to incipient speciation or intraspecific diversity would provide a better understanding of the role of hyper-variable regions in long-term evolutionary processes.

In closing, our analyses broaden the metabolic diversity known from *Pelagibacteraceae* genomes and suggest that metabolic versatility has contributed to SAR11’s success in the global ocean. While metabolic reconstructions are subject to uncertainties that necessitate physiological exploration with controlled experimentation in the laboratory, the metabolic analysis of high-quality genomes from cultivated isolates provides the opportunity to delineate metabolic specialization that could advance cultivation approaches and allow for more targeted media recipes (Rappé et al. 2002; Stingl et al. 2007; Song et al. 2009; Henson et al. 2016). SAR11 is estimated to oxidize between 6% to 37% of gross ocean primary productivity (White et al. 2019). Future work combining these metabolic predictions with *in situ* measures will greatly improve our understanding of how SAR11 impacts the dissolved organic matter pool and biogeochemical cycles in the ocean.

## Methods

### Pangenome analyses

Publicly-available *Pelagibacteraceae* isolate genomes downloaded from the National Center for Biotechnology Information (NCBI) or the Joint Genome Institute (JGI) were combined with genomes sequenced from newly isolated strains (Freel et al. 2024; **Supplementary file 1a**). Genome completion and contamination were examined with checkM v1.1.2 (Parks et al. 2015) and only isolates above 90% completion with less than 5% redundancy (Bowers et al. 2017) and fewer than 50 contigs were kept (**Supplementary file 1a).** High-quality *Pelagibacteraceae* genomes were used to construct a pangenome using anvi’o v8.0 (Eren et al. 2021) following previously described pipelines (Delmont and Eren 2018). Briefly, an anvi’o database was created using ‘anvi-gen-contigs-db’ and Prodigal v2.6.3 (Hyatt et al. 2010) was used to identify open reading frames (ORFs) from contigs. Single-copy core genes were identified using HMMER v3.2.1 (Eddy 2011). ORFs with associated functions were annotated from NCBI’s Clusters of Orthologous Groups (COGs; Tatusov et al. 2003) and a customized HMM database of KEGG orthologs (KOfams; Kanehisa and Goto 2000; Aramaki et al. 2020). The pangenome was created with ‘anvi-pan-genomè, which uses NCBI’s Basic Local Alignment Search Tool (BLAST; Altschul et al. 1990) to quantify the similarity between pairs of gene clusters and the Markov Cluster algorithm (MCL; Enright et al. 2002) to define homologous gene clusters with a MCL inflation parameter of 2. The pangenomes were visualized using the command ‘anvi-display-pan’ and summary tables exported using the command ‘anvi-summarize’.

Genome-wide average nucleotide identity (gANI) was estimated using FastANI in anvi’o (Jain et al. 2018). A phylogenomic tree of the *Pelagibacteraceae* was estimated using IQ-Tree v2.12 (Minh et al. 2020) using 1000 ultrafast bootstraps and model LG+F+R10 from a concatenated alignment of a custom gene set for SAR11 (SAR11_165; Freel et al. 2024).

Phylogenies were rooted and branches trimmed in R v 4.4.1 (R Core Team 2023) using treeio v1.28.0 (Wang et al. 2020) and visulized in R using phytools v 2.3.0 (Revell 2012) to examine the phylogenomic relationships between *Pelagibacteraceae* groups detected in our environmental study.

Gene clusters found in 100% of *Pelagibacteraceae* genomes were considered core while gene clusters found in only one genome were considered singletons. Gene clusters were also assessed as genus specific (shared among all representatives of a genus and not found in other genera) and multi-genus specific (shared among all representatives of two or more genera and not found among all others). Shared gene content between genera was assessed using ComplexUpset v 1.3.3 (Lex et al. 2014).

### Metagenomic read recruitment and environmental analyses

To examine the distribution of *Pelagibacteraceae* genera within the environments from where the majority were isolated, metagenomic read recruitment was conducted using surface ocean metagenomes from 10 stations within and adjacent to Kānʻeohe Bay, Hawaiʻi (**Fig. 1**) collected as part of the Kāneʻohe Bay Time-series (KByT; PRJNA971314; Tucker et al. 2021, 2024). In addition, metagenomes collected in the surrounding North Pacific Subtropical Gyre at Station ALOHA, a sampling location within the Hawaii Ocean Time-series (**Fig. 1; Supplementary file 1c**; PRJNA352737; Mende et al. 2017) were also used. Metagenomes were competitively mapped with Bowtie2 v 2.3.5 (Langmead and Salzberg 2012) to the anvi’o contig database of *Pelagibacteraceae* isolate genomes. The ‘anvi-profile’ function stored coverage and detection statistics of each *Pelagibacteraceae* genome found in the KByT and Station ALOHA metagenomic samples and the ‘anvi-meta-pan-genome’ function (Delmont and Eren 2018) was used to bring together the pangenomic information with the read recruitment data.

To evaluate the distribution of individual genomes, a detection metric-the proportion of the nucleotides in a given sequence that are covered by at least one short read-was used to define whether a population was detected in a metagenomic sample. A detection value of at least 0.25 was used as criteria to eliminate false positives that could arise if an isolate genome was falsely found within a sample (Utter et al. 2020).

The average depth of coverage excluding nucleotide positions with coverages in the 1^st^ and 4^th^ quartiles (mean coverage Q2Q3) was mapped for each genome in each sample. To avoid biased estimates of coverage that can occur due to highly recruiting accessory genes, read recruitment was analyzed using only single-copy genes core to each genus. Using a custom script in R, gene clusters that were found within all genomes of the genus (e.g. core), but found only in a single copy within each genome were identified. Then the mean coverage Q2Q3 read recruitment data per gene were then subsetted for only single copy core genes. The mean coverage Q2Q3 of each single-copy core gene was summed per genome per sample and then this value was normalized per genome per sample, based on whether the genome was detected at >0.25 (Utter et al. 2020). Next, when evaluating read recruitment at the genus-level, the read recruitment was summed for all genomes in a genus per sample. These outputs were then divided across the total *Pelagibacteraceae* read recruitment for the sample to yield a relative estimate of each genome or genus within a sample. For the genus with only a single genome representative (*Semelpelagibacter*), the single-copy core genes shared with the HIMB1483 genome and nearest neighbor, genus *Lacunapelagibacter*, were used.

K-means clustering analysis of a scaled matrix of biogeochemical conditions was used to characterize the environmental background from which the metagenomic data were derived.

Metadata from Station ALOHA were downloaded from https://hahana.soest.hawaii.edu/hot/hot-dogs/ (accessed 11 Oct 2021). All KByT (n=158) and Station ALOHA (n=34) surface seawater samples (depth of <30m) with data for flow cytometrically-determined cellular abundance, chlorophyll *a* concentrations, silicate concentrations, temperature, pH, and salinity were included in the analyses (**Supplementary file 1c**). The number of clusters were estimated using the ‘kmeansruns’ command in the R fpc v 2.2.13 package (Hennig 2024) using a Calinski Harabasz index. Principal Component Analysis (PCA) was conducted using the ‘prcomp’ function in R v 4.4.1 (R Core Team 2023) and visualized using ‘ggbiplot’ in the ggplot2 v 3.5.1 (Wickham 2016).

The sampling locations of the Kāneʻohe Bay Time-series and Station ALOHA were mapped using R with ‘geom_sf’ from ggplot2 with geospatial data of the main Hawaiian Islands (USGS Digital Line Graphs). To visualize *Prochlorococcus* cellular abundance across the Kāneʻohe Bay Time-series, ‘mba.surf’ from MBA v 0.1.2 (Finley et al. 2017) was used to interpolate data over the KByT stations.

### Functional inferences from *Pelagibacteraceae* genera

Functional enrichment analysis was performed in anvi’o and has been described previously (Shaiber et al. 2020). Briefly, the ‘anvi-compute-functional-enrichment’ command, as well as the ‘anvi-script-gen-function-matrix-across-genomes’, was used to assess the enrichment of Clusters of Orthologous Groups (COG), KEGG orthologs (KOs), and KEGG modules across genomes and genus affiliations. The degree of completeness of individual KEGG modules (Kanehisa et al. 2014, 2017) in the genomes and genera was evaluated using ‘anvi-estimate-metabolism’ (Veseli et al. 2023), prior to ‘anvi-compute-functional-enrichment’. The functional pangenome was visualized using ‘anvi-display-functions’. Central and alternative carbon metabolisms, amino acid, vitamin, and cofactor biosynthesis, and genes to cope with nutrient, osmotic, and oxidative stress were examined across the *Pelagibacteraceae* isolate genomes. The distribution of functional genes were evaluated for each genus as follows: core genes found in all genomes of a genus, intermediate genes found in 15-99% of genomes, and rare genes found in less than 15%. Heatmaps of the metabolic gene distribution across genera were made in the R package pheatmap v 1.0.12 (Kolde 2019).

### Selection within *Pelagibacteraceae* genes

To evaluate the selective pressures on individual metabolic genes of interest, the proportion of non-synonymous to synonymous (pN/pS) sites per gene across genomes was evaluated. The pipeline followed those developed in Kiefl et al., (2023). Briefly, contig databases from the type genomes of each of the *Pelagibacteraceae* genera (Freel et al. 2024) were first functionally annotated in anvi’o (see above) and metagenomic reads from deeply sequenced metagenomes from either coastal Kāneʻohe Bay or the adjacent offshore were non-competitively recruited using Bowtie2 v 2.3.5 (Langmead and Salzberg 2012; **Supplementary file 1c**).

Next, thèanvi-profile’ function with the flag ‘--profile-SCVs’ was used to characterize single codon variants across the read recruitment data and then ‘anvi-gen-variability-profile’ was used to export the single codon variants per gene per sample using the flags--engine CDN --include-site-pN/pS --kiefl-mode (Kiefl et al. 2023). Finally, pN/pS values were reported per gene with at least 20x coverage in each sample using ‘anvi-get-pn-ps-ratio’ with the-m 20 flag. Per gene coverage statistics were exported using ‘anvi-summarize’ with the ‘--init-gene-coverages’ flag. Functional annotations were exported using ‘anvi-export-functions’.

To evaluate whether a single genome representative contained similar pN/pS gene values as other genomes within a given genus, these steps were repeated with all representatives of the genus sharing the lowest minimum gANI (among the genera that were abundant in the tropical Pacific): genus *Coralipelagibacter*. The exported data files were brought into R, where variation in pN/pS per sample, per genome, and across genomes of the same genus was evaluated.

Relationships between pN/pS and coverage were examined and genes per genome were ranked from lowest pN/pS value to highest and visualized in R using ggplot2 v 3.5.1 (Wickham 2016). Figures were made in R v 4.4.1 (R Core Team 2023) or anvi’o (Eren et al. 2021) and further edited with Inkscape (https://inkscape.org/).

## Data availability

Accession numbers for genome sequences used to conduct pangenomic analyses and read recruitment in this study are found in **Supplementary file 1a**. Short-read metagenome accession IDs and environmental data used in this study are provided in **Supplementary file 1c**. Long-read metagenomes from coastal Kāneʻohe Bay are available under NCBI Project PRJNA1201851.

## Code availability

Code to run analyses and create figures can be found at https://github.com/tucker4/SAR11_metabolism.

## Acknowledgements

We thank Oscar Ramfelt and Evan B. Freel for their insights on computational analyses. This research was funded by the Simons Postdoctoral Fellowships in Marine Microbial Ecology (LS-FMME-00989028), the NOAA Margaret A. Davidson Fellowship (No. NA20NOS4200123), the National Science Foundation Graduate Research Fellowship Program under Grant No. 1842402, the Colonel Willys E. Lord, DVM and Sadina L. Lord Scholarship, the ARCS Foundation George Orton and Mona Marie Elmore Award in Marine Biology, the Research Corporation of the University of Hawaiʻi Graduate Research Fellowship, the Ecology, Evolution and Conservation Biology (EECB) Hampton and Meredith Carson Fellowship, and the Center for Microbial Analysis through Island Knowledge & Investigation Research Grant to SJ Tucker, and NSF grants OCE-1538628 and OCE- 2149128 to MS Rappé. AME acknowledges support from the Center for Chemical Currencies of a Microbial Planet (C-CoMP) (NSF Award OCE-2019589, C-CoMP publication #065), and Simons Foundation (grant #687269).

## Extended Data Figures

**Extended Data Fig. 1.**
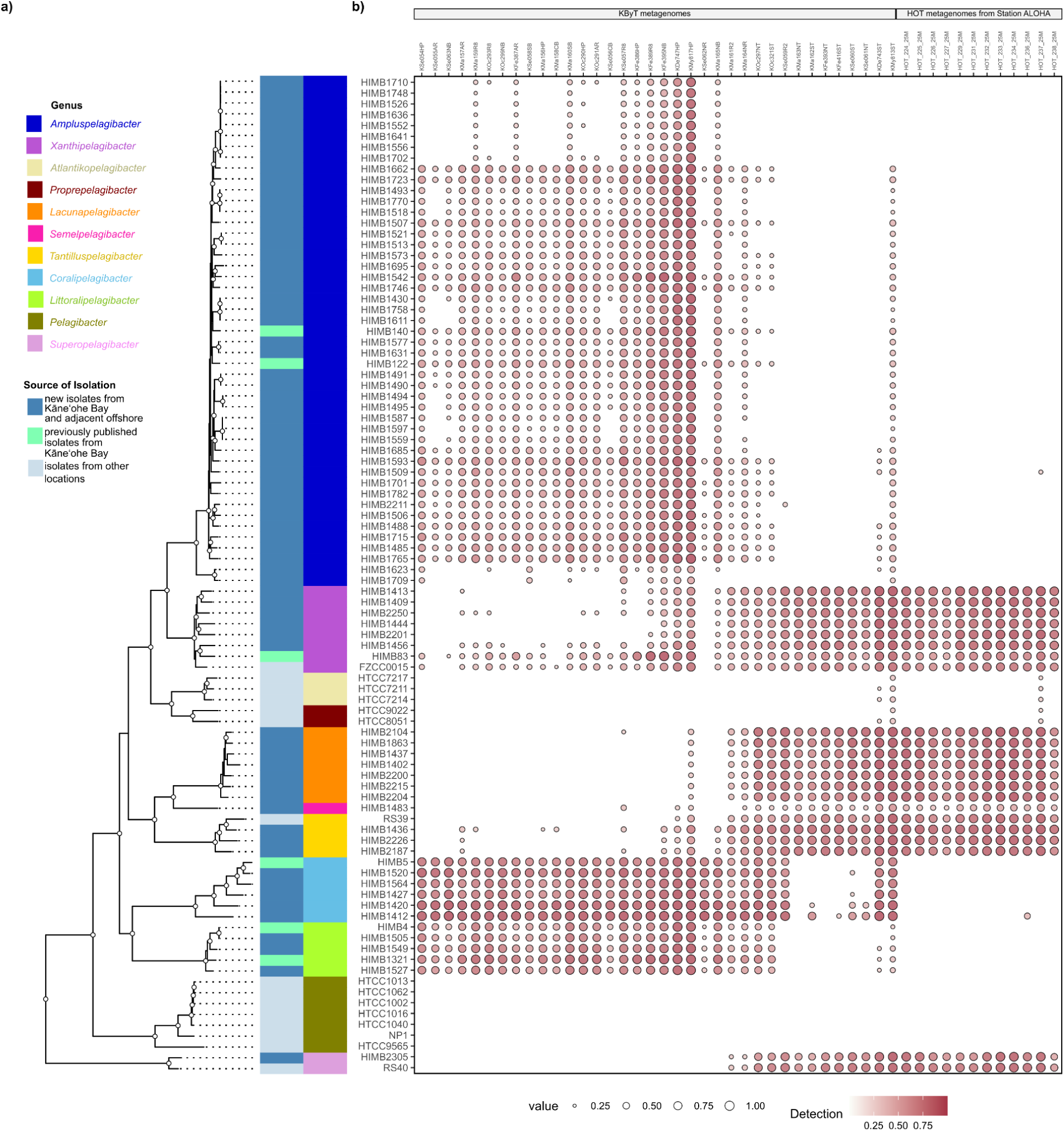
Phylogenomic position of *Pelagibacteraceae* isolate genomes and detection in the tropical Pacific. **a)** Phylogenomic tree showing 11 monophyletic clusters within the *Pelagibacteraceae*, their genus assignments, and source of isolation. The majority of *Pelagibacteraceae* isolate genomes were isolated from the KByT system. Circles at nodes indicate ultrafast bootstrap support values ≥90% from 1000 replicates. **b)** Detection of *Pelagibacteraceae* genomes in metagenomic samples from KByT and HOT at Station ALOHA. *Pelagibacteraceae* genera that contained KByT isolate genomes were commonly detected within the KByT system and sometimes at Station ALOHA. The three genera not containing KByT isolates were rarely detected in tropical Pacific, with little detection in samples collected in the KByT system or at Station ALOHA. The order of metagenomes presented follows the order in **Supplementary file 1d**.

**Extended Data Fig. 2.**
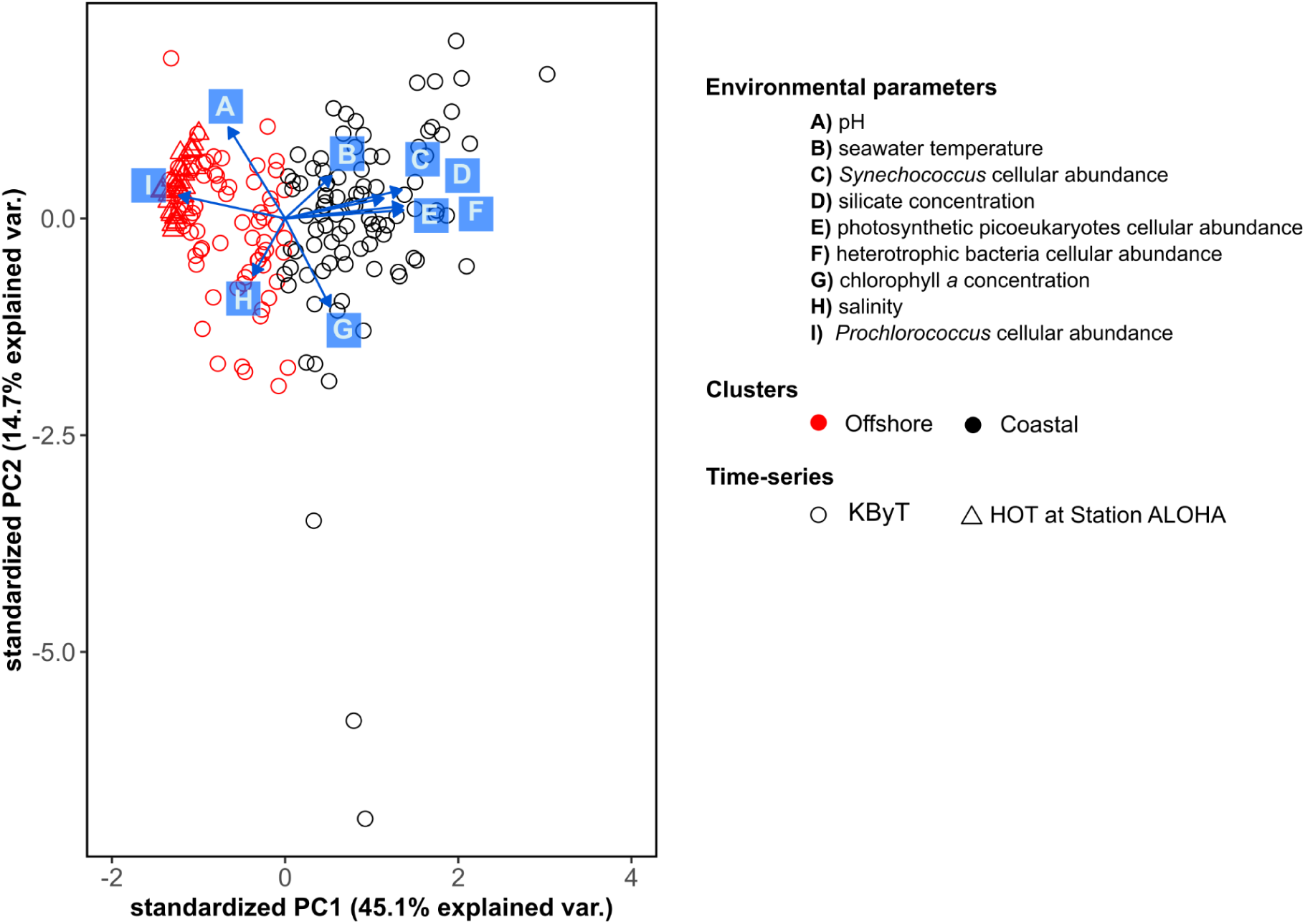
Surface ocean samples from KByT and HOT at Station ALOHA grouped into two clusters. K-means analyses of environmental parameters from 192 surface ocean samples from KByT and HOT at Station ALOHA grouped into two clusters, herein referred to as coastal and offshore. The underlying environmental covariates from the clustering analysis were mapped using Principal Components Analysis (PCA), which explained 45.1% and 14.7% of the variation across PC1 and PC2, respectively.

**Extended Data Fig. 3.**
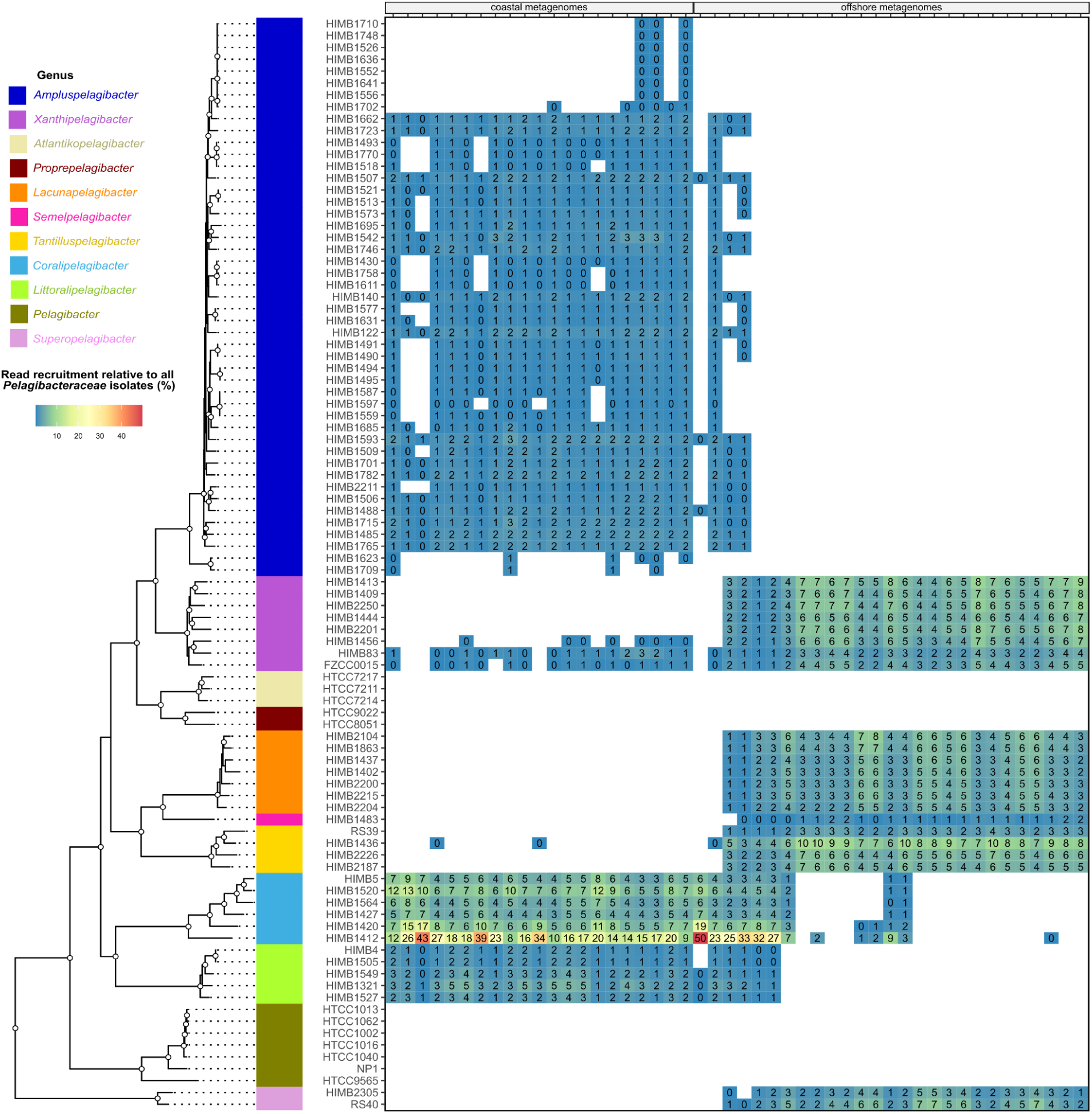
*Pelagibacteraceae* genomes sharing the same genus show similar biogeographies. Relative read recruitment of each isolate genome across the coastal and offshore environment shows that genomes belonging to the same genus share similar distribution patterns. Genera tend to be either highly prevalent in coastal samples or in the offshore samples, but not both. Data for genomes that are not detected at >0.25 or have a relative recruitment less than 0.25% are not shown. The order of metagenomes presented follows the order in **Supplementary file 1d**.

**Extended Data Fig. 4.**
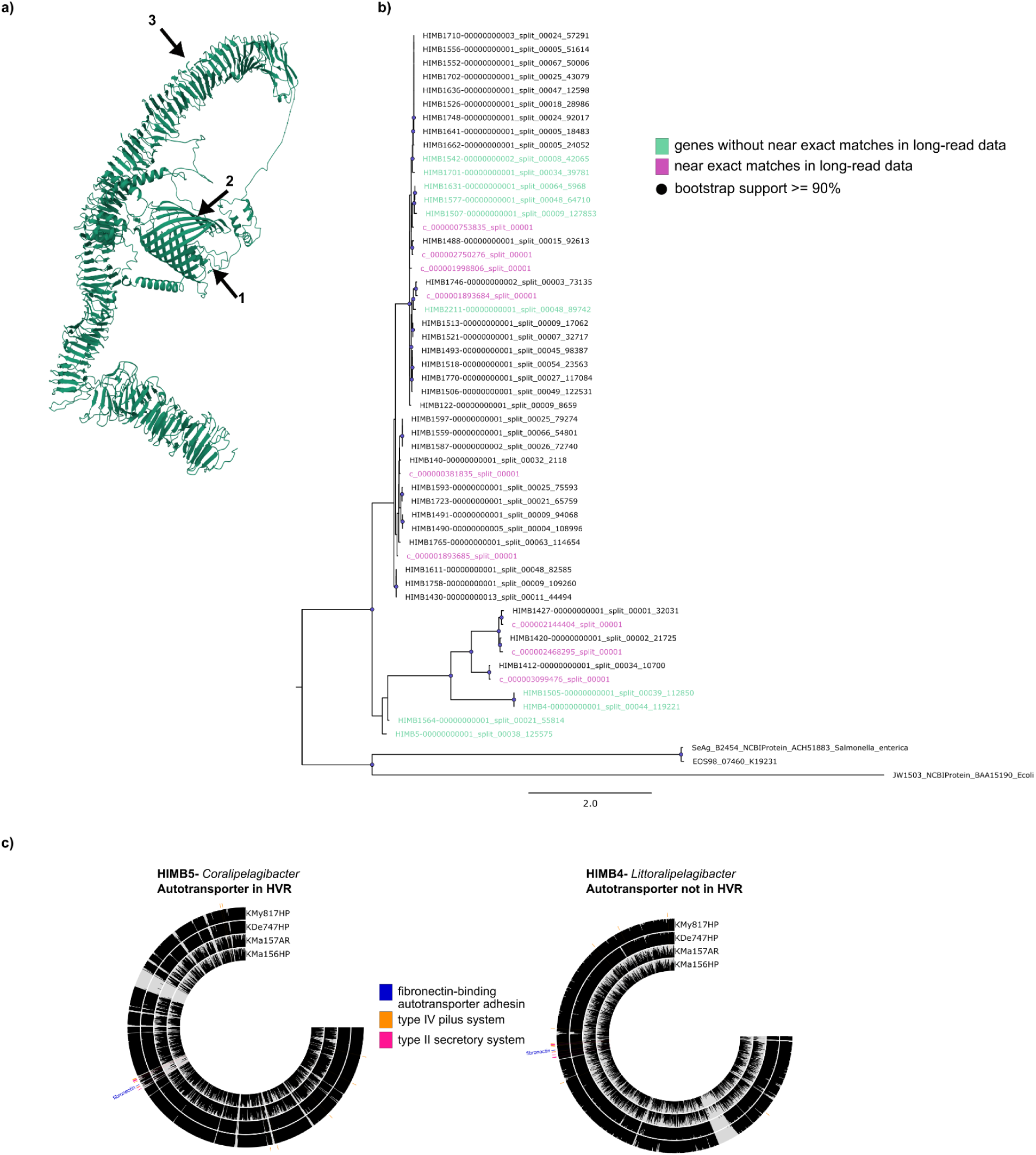
Protein structure models and homolog searches with long-read metagenomes supports autotransporter adhesion genes in coastal *Pelagibacteraceae*. **a)** A protein structure model of a gene annotated as an autotransporter adhesin from KByT long-read metagenomes that shares close sequence similarity to gene sequences in *Pelagibacteraceae* genomes has characteristics expected of autotransporter proteins: 1) α-helical linker, 2) β-barrel translocator domain, and 3) passenger domains containing β helices (Leyton et al., 2012). **b)** Homolog searches to autotransporter adhesin genes in long-read metagenomes recovered nine non-redundant gene sequences that were near exact matches to most gene sequences in *Pelagibacteraceae* genomes (33 of 43). **c)** Patterns of detection of short-read metagenomic read recruitment data across genes in two genomes of *Pelagibacteraceae* show that autotransporter adhesion genes are sometimes located in hypervariable regions and sometimes not. Genes for each *Pelagibacteraceae* genome are ordered by genome synteny with detection values of read recruitment data per gene shown as a bar per metagenomic sample. Areas of low detection are indicative of hypervariable regions (HVRs), and the autotransporter adhesin genes were located in HVRs within the HIMB5 genome, but not within HIMB4 genome. The autotransport adhesion genes were always positioned next to genes involved in type IV pilus systems and/or type II secretory systems.

**Extended Data Fig. 5.**
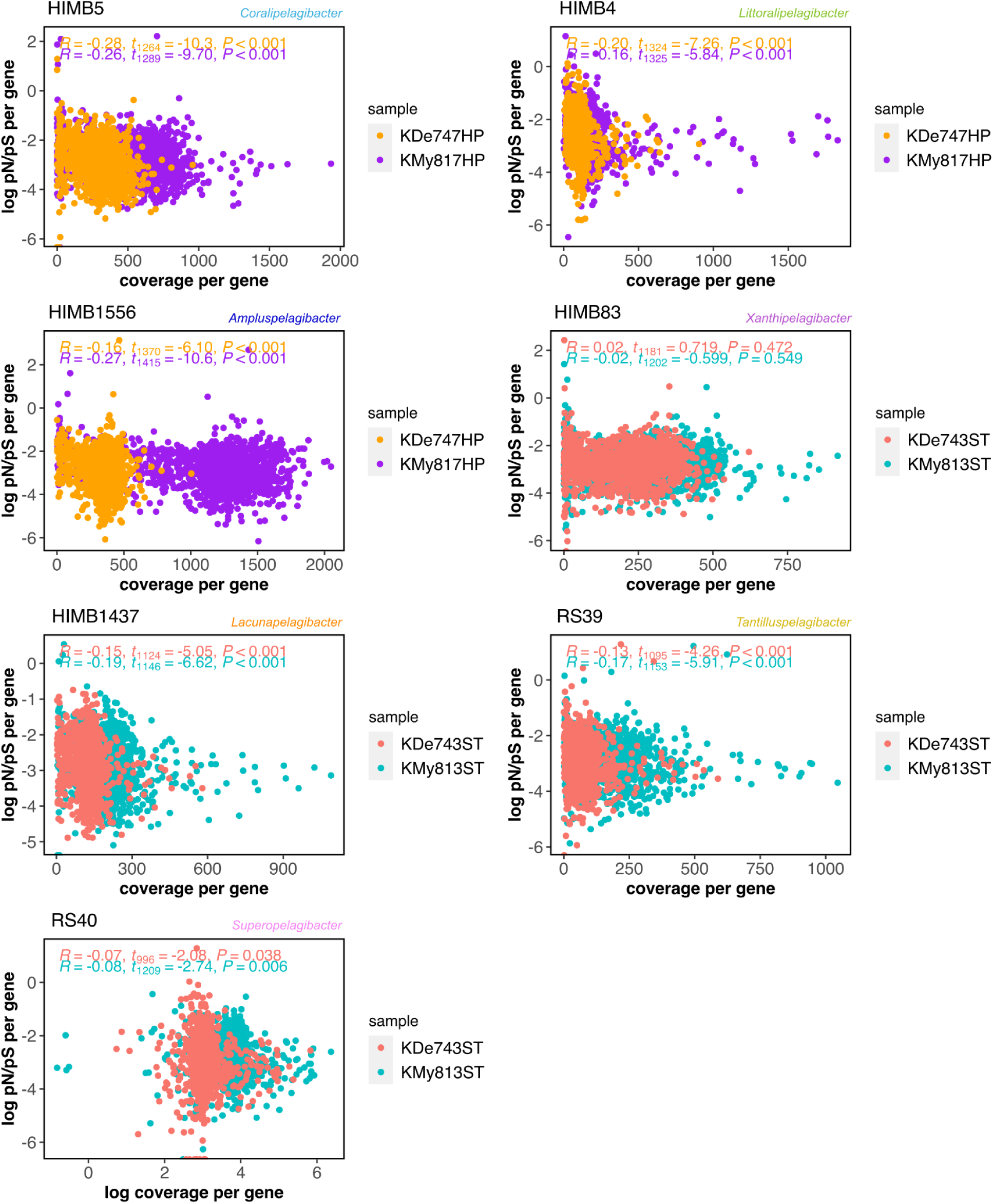
Relationship between pN/pS values and gene coverages in *Pelagibacteraceae* genomes. Reads from deeply sequenced metagenomes collected at coastal (KDe747HP, KMy817HP) and offshore (KDe743ST, KMy813ST) Kāneʻohe Bay Time-series (KByT) stations were recruited to type genomes for each of the seven *Pelagibacteraceae* genera abundant in the KByT system. The relationship between coverage per gene and pN/pS values per gene were compared for each of the genomes, showing that gene coverage did not correlate with pN/pS value. Data were normalized with log transformations. pN/pS: proportion of non-synonymous to synonymous.

**Extended Data Fig. 6.**
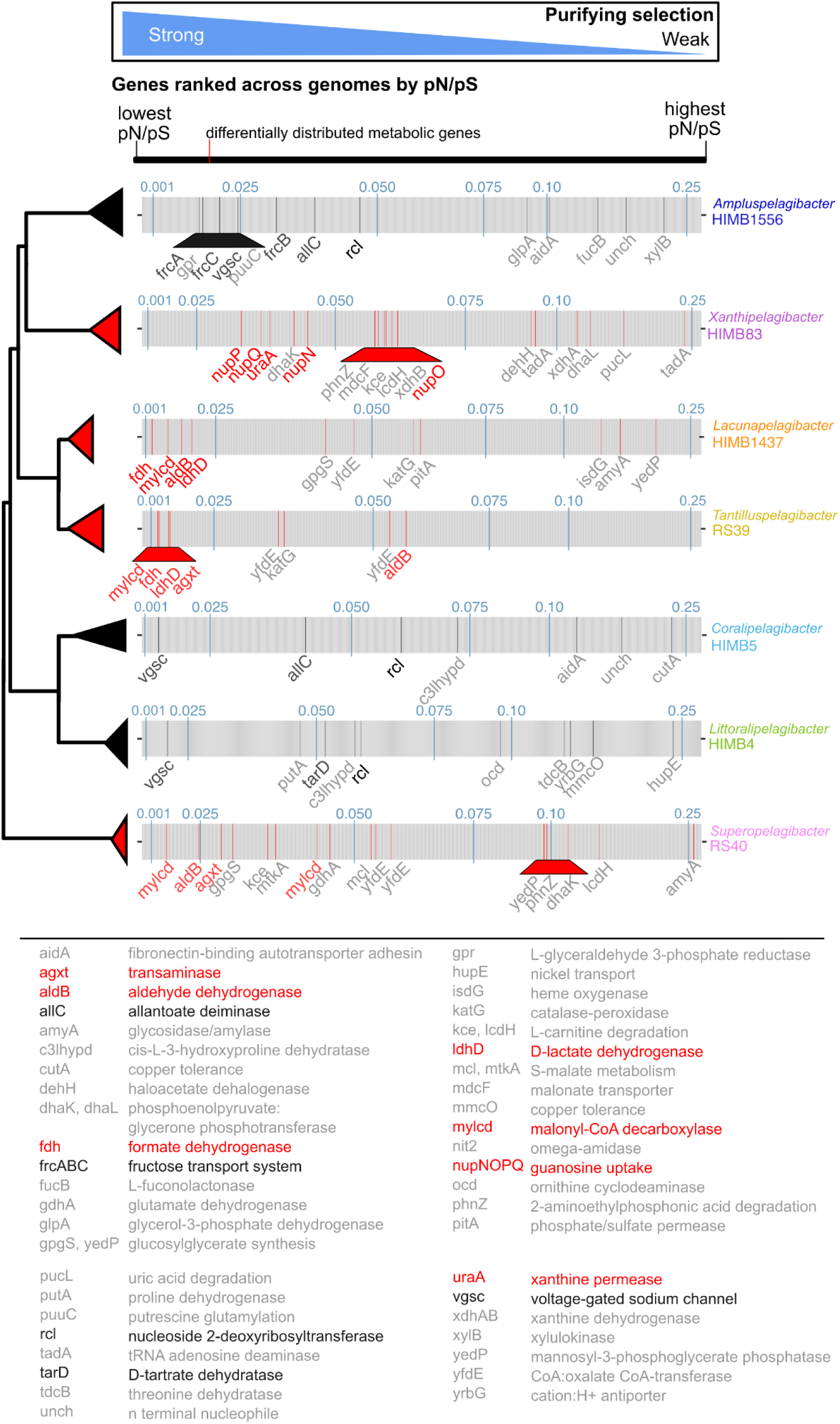
Selective pressures on habitat-specific genes in *Pelagibacteraceae*. All genes in a genome are ordered by lowest pN/pS value (high purifying selection) to highest pN/pS values (low purifying selection). Genes that are differentially distributed between offshore and coastal *Pelagibacteraceae* genera in Fig. 4 are colored red (offshore) or black (coastal) across the genomes. Gene labels that are colored in red or black are those that were highlighted in the main text and shown in Fig. 5. pN/pS values of 0.01, 0.025, 0.05, 0.075, 0.1, and 0.25 are shown in blue. Genomes are ordered by the phylogenomic relationships. pN/pS: proportion of non-synonymous to synonymous.

**Extended Data Fig. 7.**
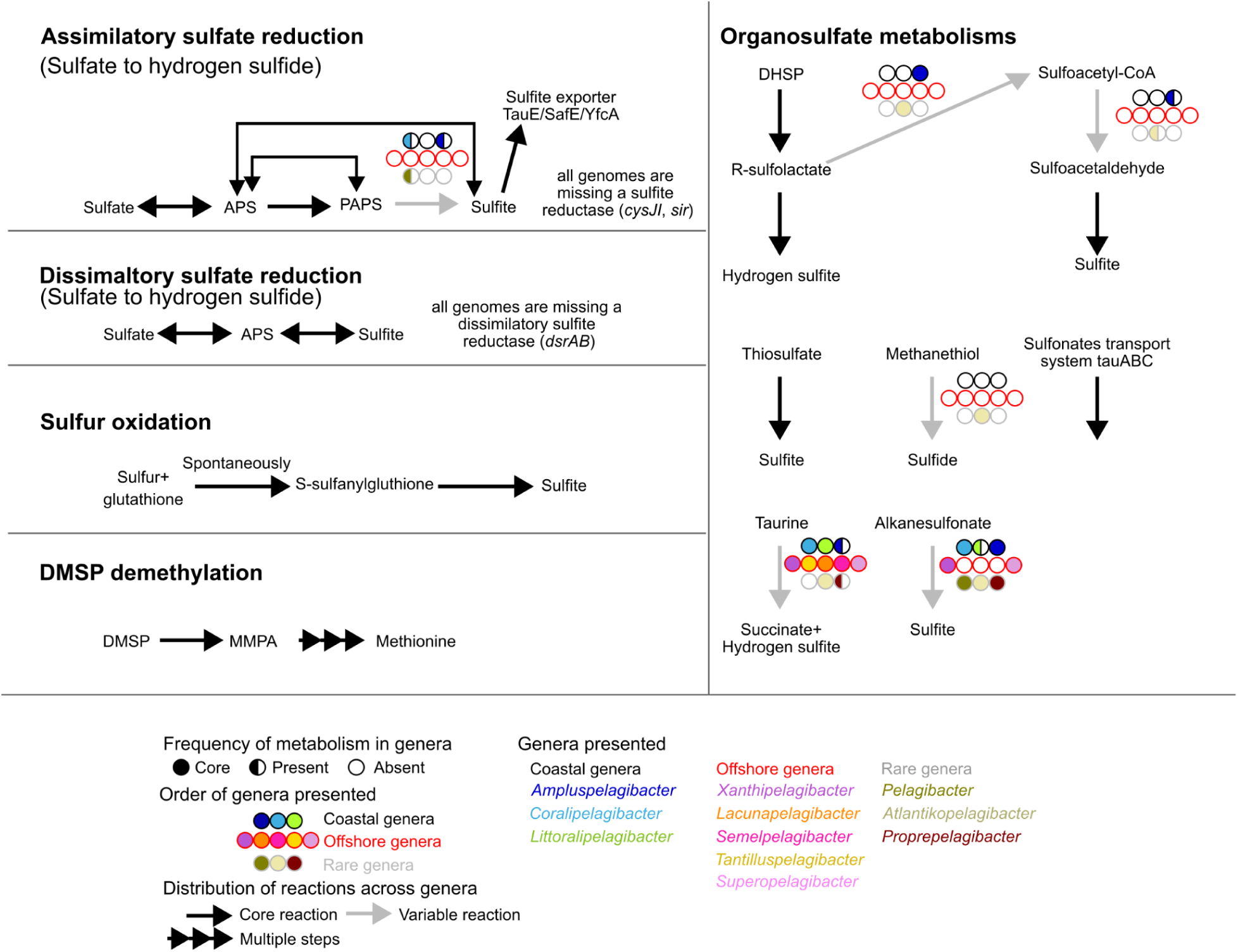
Sulfur metabolisms within *Pelagibacteraceae*. *Pelagibacteraceae* genomes have incomplete dissimilatory and assimilatory sulfur reduction pathways and the capacity to utilize a wide variety of organosulfate compounds. APS: adenylyl sulfate, PAPS: 3’-phosphoadenylyl sulfate, DHSP: 2,3-dihydroxypropane-1-sulfonate, DMSP: dimethylsulfoniopropionate, MMPA: methylmercaptopropionate.

## Supplementary Information

All supplementary infomation text and data files are available on Figshare at: https://doi.org/10.6084/m9.figshare.28087925.v3.

